# LIS1 RNA-binding orchestrates the mechanosensitive properties of embryonic stem cells in AGO2-dependent and independent ways

**DOI:** 10.1101/2022.03.08.483407

**Authors:** Aditya Kshirsagar, Anna Gorelik, Tsviya Olender, Tamar Sapir, Daisuke Tsuboi, Irit Rosenhek-Goldian, Sergey Malitsky, Maxim Itkin, Amir Argoetti, Yael Mandel-Gutfreund, Sidney R. Cohen, Jacob Hanna, Igor Ulitsky, Kozo Kaibuchi, Orly Reiner

## Abstract

*Lissencephaly-1* (*LIS1*) is associated with neurodevelopmental diseases and is known to regulate the activity of the molecular motor cytoplasmic dynein. Here we show that LIS1 is essential for the viability of mouse embryonic stem cells (mESCs), and it regulates the physical properties of these cells. LIS1 dosage substantially affects gene expression, and we uncovered an unexpected interaction of LIS1 with RNA and RNA-binding proteins, most prominently the Argonaute complex. We demonstrate that LIS1 overexpression partially rescued the expression of extracellular matrix (ECM) and mechanosensitive genes conferring stiffness to Argonaute null mESCs. Collectively, our data transforms the current perspective on the roles of LIS1 in post- transcriptional regulation underlying development and mechanosensitive processes.

## Introduction

Lissencephaly-1 (*LIS1*) was the first gene to be identified as involved in a neuronal migration disorder^1^. Proper expression levels of the LIS1 protein are critical for both mouse and human brain development, with either decreased or increased expression affecting the developmental process ^2–4^. The elimination of LIS1 is lethal during early development in mice and flies ^2, 4, 5^. LIS1 is known to play a critical role in both neuronal and hematopoietic stem cells ^6–14^. To date, these crucial roles of the LIS1 protein have been mainly attributed to its physical interaction with cytoplasmic dynein, which has been conserved throughout evolution ^15–17^. The direct binding of LIS1to dynein and additional accessory proteins results in conformational changes and modified mechanochemical properties of the molecular motor ^18–21^. The interaction between LIS1 and cytoplasmic dynein impacts the many processes in which cytoplasmic dynein is involved, such as mitosis, interkinetic nuclear motility, neuronal migration, intracellular transport, and neuronal degeneration ^4, 10, 13, 16, 22–25^.

LIS1 has also been implicated in additional activities unrelated to its interactions with the molecular motor. LIS1 affects the cytoskeleton through modulation of microtubules and the actin mesh ^22, 26–29^. LIS1 localizes in the nucleus, where it interacts and affects the activity of MeCP2 ^30^. In human embryonic and neuronal stem cells, LIS1 affects gene expression and the physical properties of the colonies ^10^.

Here we investigate the dosage-related roles of LIS1 in embryonic stem cells using a multidisciplinary approach and detect novel and unexpected functions for LIS1 in post-transcriptional regulation and affecting the physical properties of mESCs. We found that LIS1 binds RNA and interacts with numerous proteins, many of which are RNA-binding proteins (RBPs), including some belonging to the Argonaute complex. LIS1 is mainly bound to the nascent RNA of protein-coding genes, and the number of LIS1- binding sites within introns were negatively correlated with intron splicing efficiency. A different outcome was noted when LIS1 bound to microRNA (miRs), which coincided with their increased expression. Overexpression of LIS1 in the absence of AGO1-4 enabled low but significant expression of a subset of miRs. Whereas AGO1-4 KO cells were soft, LIS1 overexpression changed the expression of genes related to the extracellular matrix and mechanosensitivity. It resulted in a robust increase in the stiffness of mESCs lacking Argonaute proteins. Collectively, these findings necessitate the need to re-evaluate the roles of LIS1 during development.

## Results

### Dosage sensitive effects of LIS1 in pluripotency networks

*Lis1-/-* mice are early embryonic lethal ^2, 4, 5^, therefore we examined the localization of LIS1 in E3.5 pre-implantation embryos. LIS1 was detected in the cytoplasm as well as the nucleus, where it partially colocalizes with either OCT4 or NANOG (Fig. 1a), suggesting unknown nuclear functions. The transcription factors OCT4 and NANOG regulate the expression of the dynamic transcription network and are required for the pluripotency and proliferation of embryonic stem cells (ESCs) ^31–34^. So far, it has not been established whether *Lis1-/-* mESCs are viable. To generate null cells, *Lis1 floxed/-:Cre-ERT2 (Lis1 F/-:ERT2),* mESC lines were derived from blastocysts, and the *Lis1* deletion was induced by tamoxifen (4-OHT) treatment. When *Lis1* was deleted in the presence of the standard 2i+LIF media, the cells died rapidly (Fig. 1B). We then reasoned that 5i+LIF media, modified from human naive media ^35^, may support cell viability (Extended Data Table 1). mESC colonies containing the Oct4 reporter line ^36^ appeared compact and pluripotent in this media compared to 2i+LIF, or FBS+LIF (Extended Data Fig. 1a). The necessity to remove the Wnt inhibitor (XAV939) in the 5i+LIF media was suggested due to the findings that reduced LIS1 dosage in brain organoids resulted in inhibition of the Wnt pathway ^37^, and was supported by examining gene expression (Extended Data Fig. 1b). The 5i+LIF modified media-enabled three passages of cultured *Lis1-*deleted cells, after which the cells died (Fig. 1b). These findings demonstrated that LIS1 expression is required for the viability of mESCs. The lethality of *Lis1* deletion was rescued by ectopic expression of LIS1- GFP (Fig. 1B). RNA-expression data was obtained from the different LIS1-dosage-dependent genotypes (Fig. 1c). When comparing the differentially expressed (DE) genes between the highest and the lowest LIS1 dosage (LIS1-GFP OE, floxed/- background versus floxed/- treated with tamoxifen, that is *Lis1-/-,* respectively), 2150 genes were upregulated, and 904 were downregulated (Extended Data Table 2). The DE genes came together in four k-means clusters (Fig. 1c). Analysis of enriched Gene Ontology Biological Process (GO-BP) terms showed that LIS1 dosage affects many fundamental processes, including DNA replication, mitosis, apoptosis, and autophagy. RNA-related functions such as biogenesis, splicing, and non-coding RNA processing were also enriched in this analysis. LIS1 is also involved in biosynthetic and metabolic processes, including some pertaining to nucleotides, fatty acids, and organophosphate (Fig. 1d). Metabolomic analysis of mESCs revealed changes in nucleotides such as deoxyuridine, deoxycytidine monophosphosphate, and dAMP, different levels of amino acids such as proline and aspartate, and changes in essential metabolism molecules such as nicotinamide adenine dinucleotide (NAD). The fatty acid metabolic profile showed changes in monounsaturated fatty acids such as erucic acid-like and polyunsaturated fatty acids such as eicosapentaenoic acid (EPA) and docosahexaenoic acid (DHA) (Extended Data Table 3).

**Fig. 1:**
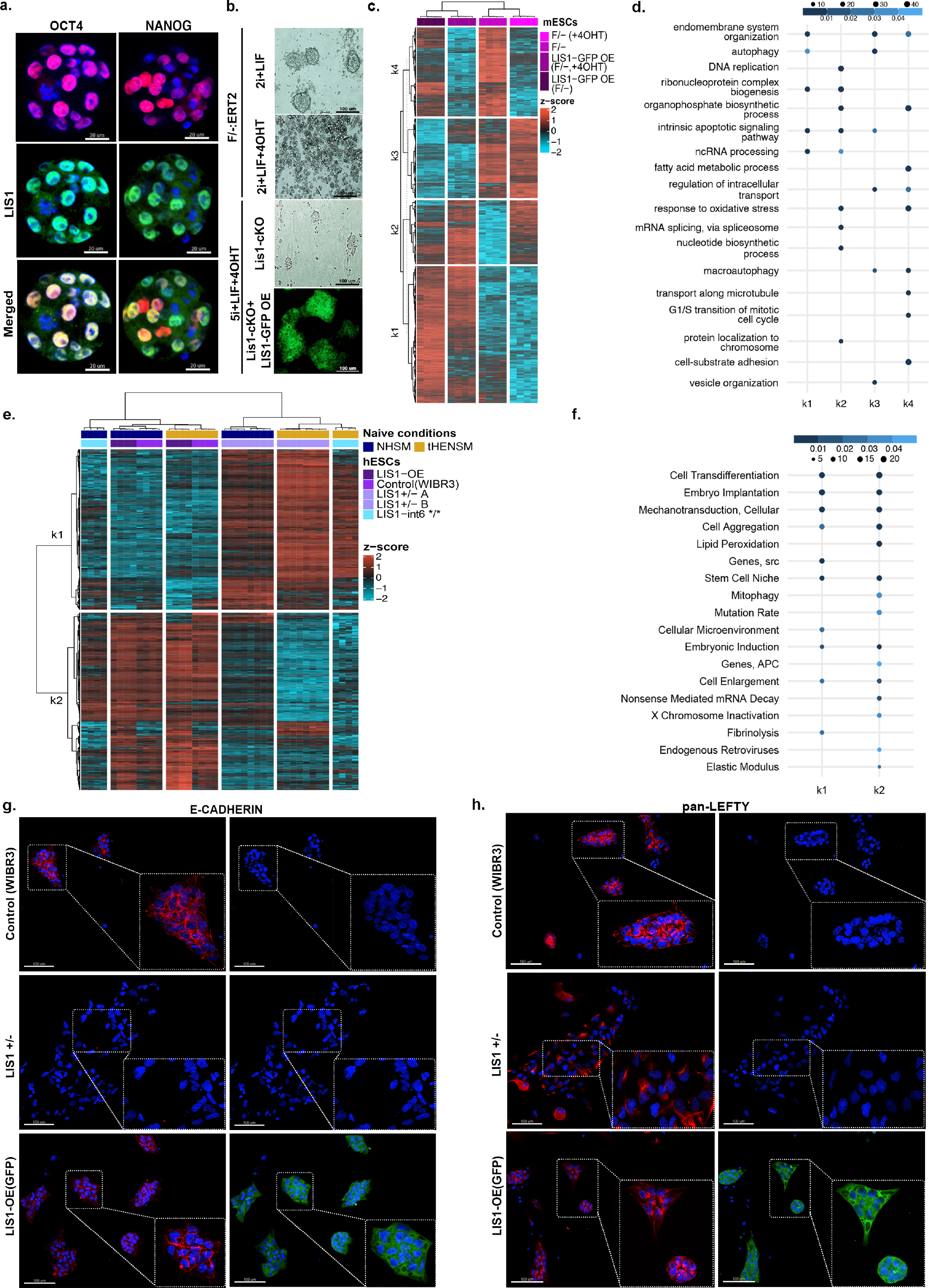
L**I**S1 **dosage affects gene expression. a.** LIS1 colocalizes with NANOG and OCT4 in the nucleus. Pre-implantation embryos from embryonic day 3.5 were immunostained with anti- NANOG, anti-OCT4, and anti-LIS1 antibodies (scale bars, 20 μm.) **a.** Phase-contrast and fluorescent images of Lis1 F/-:ERT2 mESCs cultured in 2iL media with or without 4-OHT. Lis1 F/-:ERT2 and pB-LIS1GFP overexpression mESCs treated with 4-OHT using alternative naïve (5iL) media conditions. Images are representative of at least two independent experiments (scale bars,100 μm). **c.** A heatmap of 2654 differentially expressed genes across samples with different *Lis1* gene dosage in F/-:ERT2 derived mESCs cultured in 5i+LIF media. The data are shown on a Z-score scale of the variance stabilizing transformation on normalized reads. **d.** Analysis of gene set over-representation test for four clusters obtained from the k-means clustering (k1, k2, k3, and k4) shows that different levels of LIS1 modulate the expression of genes enriched in specific Gene Ontology Biological Process (GO-BP) terms. **e.** A heatmap of 927 differentially expressed genes across samples with different *Lis1* gene dosages in hESCs (including a homozygous point mutation in *LIS1* intron 6). Samples come from two independent media conditions; human naïve, NHSM, and tHENSM. The data are shown on a Z-score scale of the variance stabilizing transformation on normalized reads. **f.** Gene set over-representation test analysis compares dose-dependent upregulated and downregulated k-means clusters (k1 and k2) in hESCs enriched for Medical Subject Headings (MeSH) terms. **g-h.** Representative immunofluorescence images of immunostainings conducted in WIBR3 (control), *LIS1* +/-, and pB-LIS1GFP overexpression (LIS1 OE) isogenic hESCs cultured in tHENSM media using anti- E-CADHERIN, and pan-LEFTY (LEFTY-A and LEFTY-B) antibodies, respectively (scale bars,100 μm. Inset represents 2.5x zoom).

Moreover, genes affected by LIS1 expression are involved in vesicle organization, intracellular transport, and cell-substrate adhesion, all of which could be associated with known LIS1 functions (Fig. 1d).

Next, we examined the role of LIS1 expression in regulating pluripotency in human ESCs. We used two media conditions^35^ and five isogenic lines; the control wild-type, two previously published *LIS1+/-* lines ^10^, LIS1 overexpression (OE), and a lissencephaly-associated intronic mutation in intron 6 affecting splicing ^38^ (*LIS1-int6*/\**) in the homozygous form, that slightly reduced LIS1 levels. The overexpression and wild-type were detected in one group and *LIS1-int6*/\** clustered with the *LIS1+/-* lines only in one growth condition (Fig. 1e). A total of 927 DE genes between the wild-type and *LIS1+/-* lines were noted (Extended Data Table 4). The selected Medical Subject Headings (MeSH) terms indicated changes in the stemness and differentiation potential, RNA, and the physical properties of the cells, such as mechanotransduction and elastic modulus (Fig. 1f). We confirmed the differential expression of a few of the genes by immunostaining. E-cadherin, with its central role in embryonic stem cell pluripotency, epithelial to mesenchymal transition (EMT), and mechanotransduction, was of particular interest ^39–41^. *LIS1+/-* cells did not express E-cadherin, though the control and the LIS1-OE cells expressed it at high levels (Fig. 1g, Extended Data Fig. 1c). LEFTY, a member of the TGF-beta family, contributes to the remodeling of the extracellular matrix and regulates actin polymerization and stiffness ^42–44^. The wild-type and LIS1-OE ESC colonies expressed LEFTY in a polarized manner, whereas there was a reduced expression in the *LIS1+/-* cells (Fig. 1h). We examined the localization and expression of β-Catenin, the primary downstream target of the Wnt pathway, in hESCs and found the most striking difference in *LIS1+/-* colonies, where the signal appeared truncated (Extended Data Fig. 1d, e). Collectively, our data demonstrate that LIS1 is expressed in the nucleus and the cytoplasm and is essential for the survival of embryonic stem cells. Although our custom-designed media-enabled three cell passages, the embryonic stem cells died. LIS1 levels affected gene expression in a dosage-specific manner and were found to be involved in multiple basic cell biological processes. We hypothesized that the known interactome of LIS1 that is composed of mainly cytoskeleton-related proteins was insufficient to explain all of these changes, and we proceeded to identify novel LIS1-interacting proteins.

### The LIS1 interactome

The known LIS1 interactome is composed of 148 proteins, many of which are involved in microtubule-based processes (BioGRID^45^ and STRING^46^ databases), and very few are nuclear proteins. Novel LIS1-interacting proteins were identified by immunoprecipitation of LIS1 from nuclear or cytoplasmic fractions of mESCs expressing graded levels of LIS1 followed by mass-spectrometry (LC-MS/MS) (Fig. 2a). A total of 726 unique interacting proteins were identified, many of which were not known to complex with LIS1 (Extended Data Table 5). More than half of these proteins (385) were designated in the RBPbase (https://rbpbase.shiny.embl.de/) as “RNA binding-GO-Mm”. Most of the LIS1 interactome was shared in the cells with different LIS1 expression levels (WT, F/-, and OE) (Extended Data Fig. 2a). The top consistent and genotype-specific GO terms related to mRNA splicing and cellular response to stress (Extended Data Fig. 2b). Many LIS1-interacting proteins belonged to the RISC complex and P-bodies, interacting with AGO2 (Fig. 2b). A second prominent group of proteins was composed of RNA-binding proteins functioning in other activities, including alternative splicing, mRNA stabilization, RNA metabolism, transcriptional and translational regulation (Fig. 2c). We found several members of the heterogeneous nuclear ribonucleoproteins (hnRNPs) ^47^, small nuclear ribonucleoproteins (snRNPs)^48^, DDX, and DHX gene families ^49^. A subset of these two groups of proteins is shown in the heatmaps, demonstrating that the protein-protein interactions are enriched in the nucleus and occur in cells with graded levels of LIS1.

**Fig. 2:**
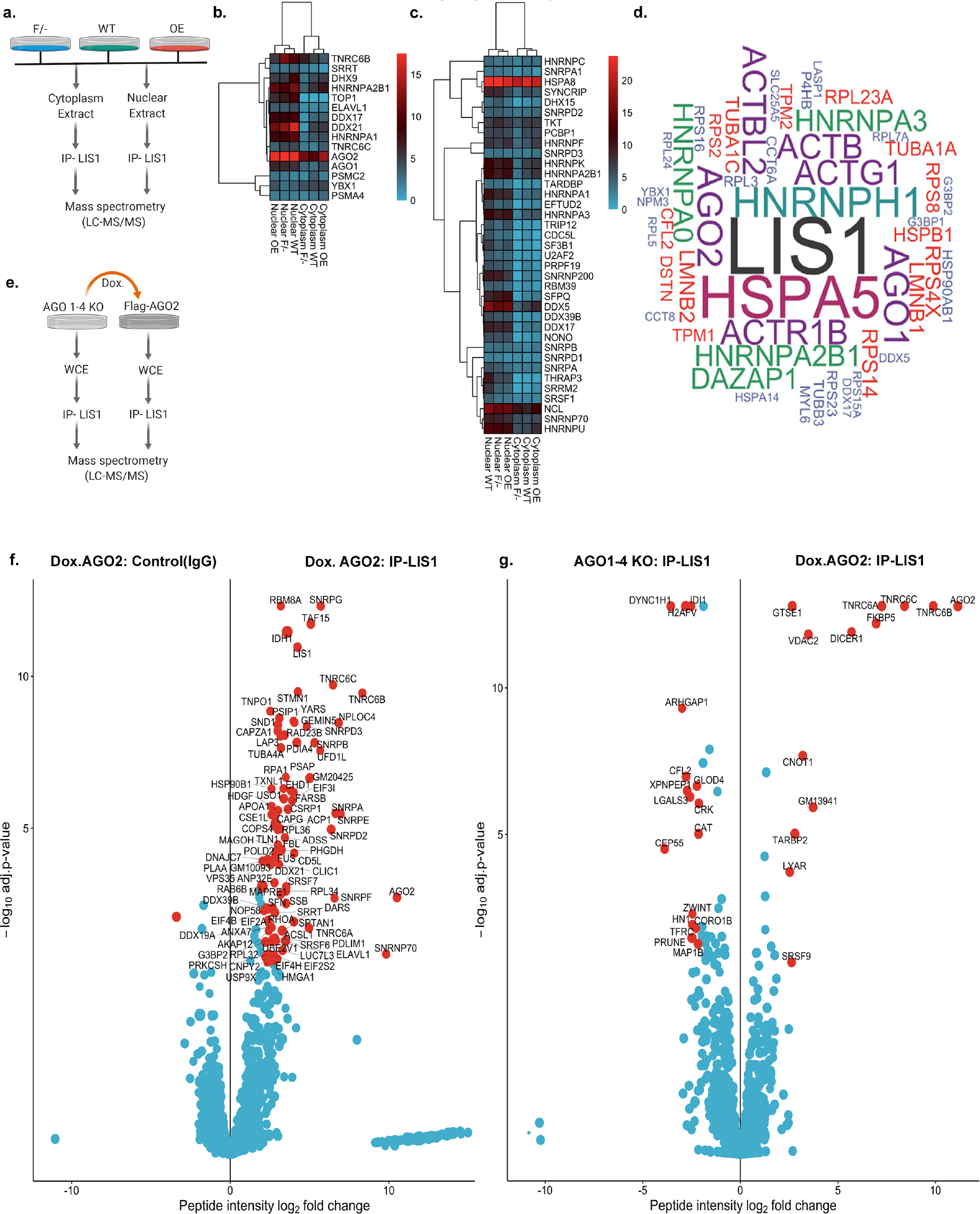
N**o**vel **LIS1-interacting proteins include a repertoire of RNA-binding proteins. a.** A schematic illustration of the strategy used to identify LIS1-interacting proteins in mESCs. The cytoplasmic and the nuclear fractions from each genotype (F/-; Floxed/-, hypomorph allele, WT; Wild Type, and OE; LIS1-dsRED overexpression) were separated. LIS1 immunoprecipitation (IP) was then performed using anti-LIS1 antibodies followed by mass spectrometry (total n=4, for each genotype). **b.** Heatmap of the RISC complex and P-body proteins derived from the mass spectrometry results. **c.** A heatmap of splicing factors and nuclear speck proteins was identified as significant LIS1 interactors. For **b.** and **c.**, the scale represents razor, and unique peptides detected averaged across replicates for each protein in each fraction per genotype. **d.** The top 50 LIS1 interactors in *Drosophila melanogaster*. The size of the word corresponds to the total number of peptides obtained in LIS1 IP for each mouse orthologue. **e.** Schematic illustration of the strategy to identify AGO2 dependent (doxycycline-inducible AGO2; Dox.AGO2^AGO1-4KO^) and independent LIS1 interacting proteins in AGO1-4 KO (AGO1/2/3/4 knockout) mESCs. LIS1 IP in combined nuclear and cytoplasmic fractions was performed in AGO1-4 KO and Dox.AGO2 followed by mass spectrometry (n=4). **f.** A volcano plot for the ratios of peptide intensitie s of proteins detected with mass spectrometry in LIS1 IP versus control (IgG, nonspecific peptides) in Dox. AGO2^AGO1-4KO^. **g.** A volcano plot for the ratios of peptide intensitie s of proteins detected in mass spectrometry with LIS1 IP in AGO1-4 KO and Dox.AGO2^AGO1-4KO^, respectively. For **f.** and **g.**, the proteins with a log2 foldchange ≥ 2 and an adjusted p-value ≤ 0.01 are highlighted red.

To assess the conservation of the identified LIS1-interactome, we compared our results with a previously published large-scale *Drosophila* interactome ^50^. A significant number of the translated *Drosophila* orthologs overlapped with the LIS1-interacting proteins in mESCs (Extended Data Fig. 2c). These included members of the AGO clade, heat-shock, chaperone proteins, and several splicing factors, suggesting that these interactions are evolutionary conserved (Fig. 2d). We further detected a significant overlap between the nuclear interactome of LIS1 and AGO2^51^ (Extended Data Fig. 2d).

To systematically identify the Argonaute-dependent LIS1 protein interactors, we used an *Ago1,2,3,4* knockout line with the doxycycline-dependent expression of human AGO2, with and without the presence of doxycycline, to perform LIS1-IP followed by mass-spectrometry ^52, 53^ (Fig. 2e and Extended Data Table 5). In the presence of AGO2, LIS1 is bound to multiple interacting proteins (including LIS1 itself) compared with control IgG, supporting the LIS1- interactome identified above (Fig. 2f). In the absence of the Argonaute proteins, LIS1 complexed with several known LIS1-interacting proteins, such as cytoplasmic dynein heavy chain, the microtubule-associated protein, MAP1B, and additional proteins, some of which have not been previously reported to interact with LIS1, such as centrosomal protein CEP55, kinetochore protein ZWINT, and actin regulator, cofilin-2 (CFL2) (Fig. 2g, left side). The induction of AGO2 expression (Dox.AGO2, Fig. 2g, right side) resulted in additional protein interactions.

These proteins included AGO2, TNRC6A-C, DICER1, FKBP5, TARBP2, and CNOT1, all of which were known to complex with AGO2. Some not previously identified as part of the AGO2- interactome include voltage-dependent anion channel pore-forming protein VDAC2, GTSE1, which may be involved in the p53-induced cell cycle arrest, and the LYAR protein involved in the processing of pre-rRNAs. We further mapped the LIS1 and AGO2 interactome to common or unique complexes (Extended Data Fig. 2e). For example, we detected “transcription repression” and “extracellular exosome” among the common GO terms.

### LIS1 is an RNA-binding protein

As we have shown that most of the LIS1 interactome is composed of RNA-binding proteins (RBPs), and previous high throughput studies identified LIS1 as an RBP,^54^, we set out to map the extent of LIS1’s interaction with RNA. Towards that end, we conducted LIS1 single-end enhanced crosslinking immunoprecipitation (seCLIP) experiments (Extended Data Fig. 3a) ^55^.

**Fig. 3:**
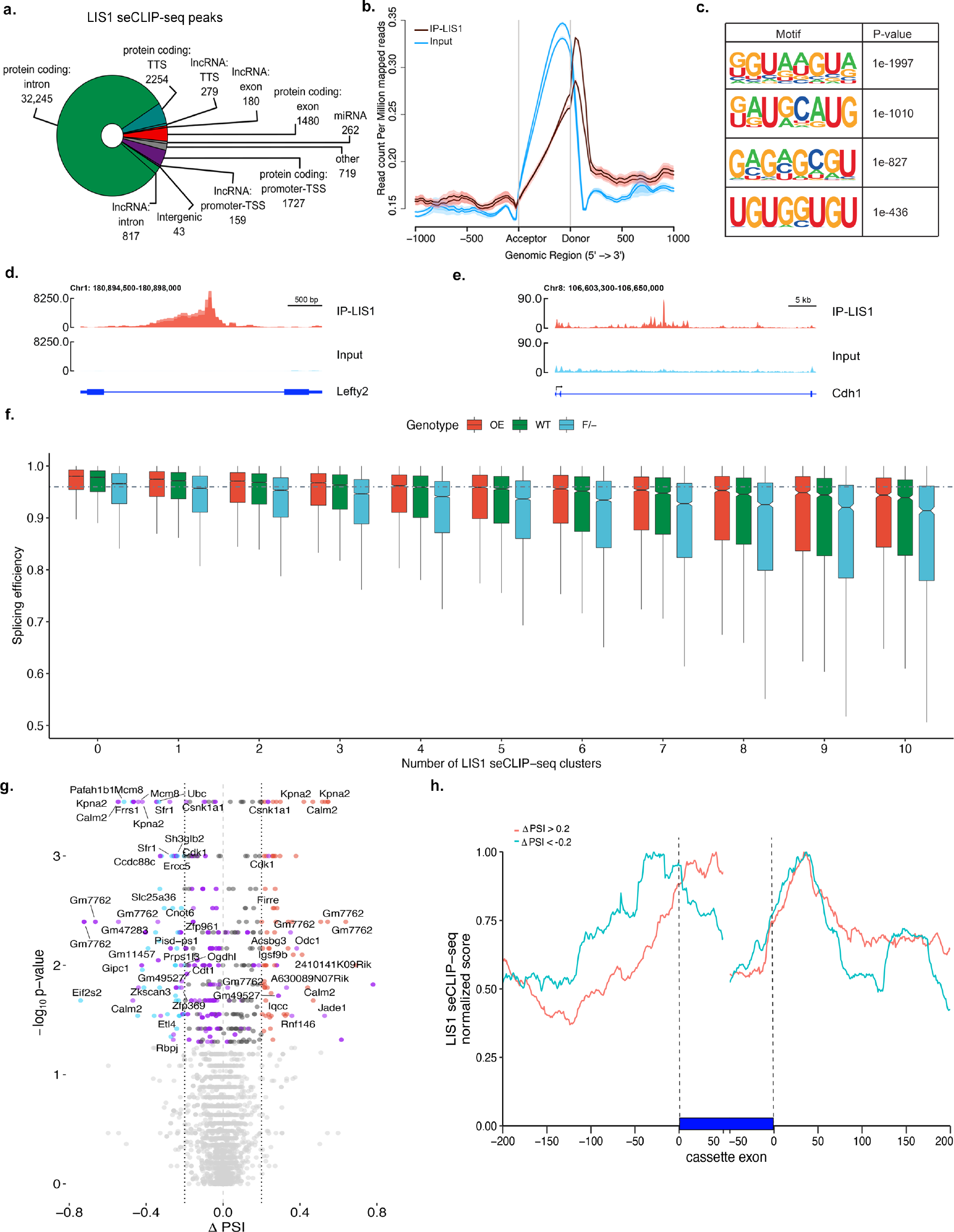
L**I**S1 **RNA-binding properties and splicing regulation. a.** LIS1 seCLIP-seq reproducible peaks (n=40165, log2 fold change ≥ 3, adjusted p-value ≤ 0.001) categorized by functional genomic regions. **b.** Coverage profile plot for the distribution of LIS1 seCLIP-seq reads across exons. Brown lines represent two individual replicates for LIS1, and blue lines indicate the input (SMI; size-matched input). **c.** Homer motifs of LIS1 seCLIP peaks. **d-e.** LIS1 eClip read coverage in *Lefty2* 1^st^ intron (D.) and E-cadherin 2^nd^ intron (E.) Red tracks- merged biological seCLIP-seq replicates (IP), Light blue tracks- input (SMI). **f**. Intron level quantification of splicing efficiency in OE (red), WT (green), and F/- (blue) with respect to the LIS1 seCLIP-seq clusters (the experiment includes n=4 RNA-seq replicates for each genotype). All the comparisons were significant across each genotype in each cluster group. The p-values are reported in extended data table 6. **g.** A volcano plot of differentially retained introns between OE and F/- was derived from the analysis of MAJIQ alternative intron usage. Significant upregulated events are in red, and (ΔPSI ≥ 0.2, p-value ≤ 0.05, Wilcoxon test) and downregulated events (ΔPSI ≤ -0.2, p-value ≤ 0.05, Wilcoxon test) are in blue. Events with retained introns and p-value ≤ 0.05 (Wilcoxon test) are highlighted in purple. The remaining events with a p-value ≤ 0.05 (Wilcoxon test) are dark grey. Light grey is events with a p-value ≥ 0.05 (Wilcoxon test). Gene symbols for the *de novo* events are shown (n=8 RNA-seq replicates for each genotype). **h.** LIS1 seCLIP signal in regions with differentially spliced events annotated cassette exons with a retained intron (ΔPSI ≥ 0.2). The red line indicates LSVs, which were found to be higher in OE, whereas the blue line indicates such events that are lower in OE (red; n= 320, blue; n= 280).

The results indicated that LIS1 binds to multiple loci (40,165 peaks), mainly found within the introns of protein-coding RNA (87%, Fig. 3a). LIS1 is preferentially bound to RNA of highly expressed genes and those with relatively large introns (Extended Data Fig. 3b,c). LIS1 seCLIP sites were more frequent in the first two introns (Extended Data Fig. 3d). Metagene plot of the data indicated that LIS1 preferentially binds to introns within close proximity to the donor splice site (Fig. 3b). A pattern of such asymmetric binding was also detected in data derived from AGO2-eCLIP experiments in both control cells and even more so in cells lacking Dicer, an essential component of the RISC complex ^56^ (Extended Data Fig. 3e,f). In addition, we noted significant two-fold enrichment of the U1 snRNP binding site in the peaks, suggesting the LIS1 may be involved in splicing regulation. LIS1-binding consensus sequences in the eCLIP peaks were GU-rich, as could be expected from intronic- or close to donor splice site sequences (Fig. 3c). Two DE genes that contain LIS1 binding sites in their introns are *Lefty2* (Fig. 3d) and the gene encoding E-cadherin, *Cdh1* (Fig. 3E), which is not expressed in the *LIS1 -/+* hESCs (Fig. 1g). Considering the preferential binding of LIS1 in proximity to donor splice sites, we tested whether LIS1 RNA-binding affected splicing. We detected a negative correlation between the number of LIS1 seCLIP sites and splicing efficiency in wild-type RNAseq data (Fig. 3f, green, Bulk RNA-Seq data in Extended Data Table 6a). When LIS1 is decreased (as in the case of F/- cells, light blue), the splicing efficiency is further reduced progressively. Conversely, the splicing efficiency was slightly but significantly increased upon LIS1 OE (Fig. 3f, red, statistics of all figures is in Extended Data Table 7a). We then examined local splicing variations (LSVs) by contrasting the LIS1 OE versus the F/- RNAseq data. Most LSVs were annotated, yet many were novel (Extended Data Fig. 4a,b). Top Reactome terms included RHO GTPase cycle, organelle biogenesis, maintenance, centrosomes, and cilia (Extended Data Fig. 4c). Examples for multiple LSVs are shown for the *Meg3* and *Rian* loci (Extended Data Fig. 4d,e). Retained intron events were plotted, showing more events occurring in the F/- (Fig. 3g). The position of LIS1 seCLIP sites in relation to cassette exons was plotted (Extended Data Table 8, Fig. 3h). While a clear peak is seen next to the donor sequence (as depicted in Fig. 3b), we noted a difference between LSVs that were higher in OE, where the peak is in the exon, and those that were lower in the OE, where the peak was in the intron close to the acceptor site (Fig. 3h). Part of the LIS1-bound genes was also DE (Extended Data Fig. 5a). In addition, for a subset of LIS1 bound DE genes exhibited differential accessibility as detected by Assay for Transposase-Accessible Chromatin using sequencing (ATAC)-seq in the different lines, including the *Meg3-Mirg* locus (Extended Data Fig. 5b) (details in methods, statistics in Extended Data Table 9). Our data indicate that LIS1 is an RBP that preferentially binds intronic sequences of highly expressed protein-coding genes near the splicing donor site. In addition, the increase in seCLIP LIS1 binding sites within a gene is negatively correlated with splicing efficiency, and increased expression of LIS1 improves the efficacy of RNA splicing.

**Fig. 4:**
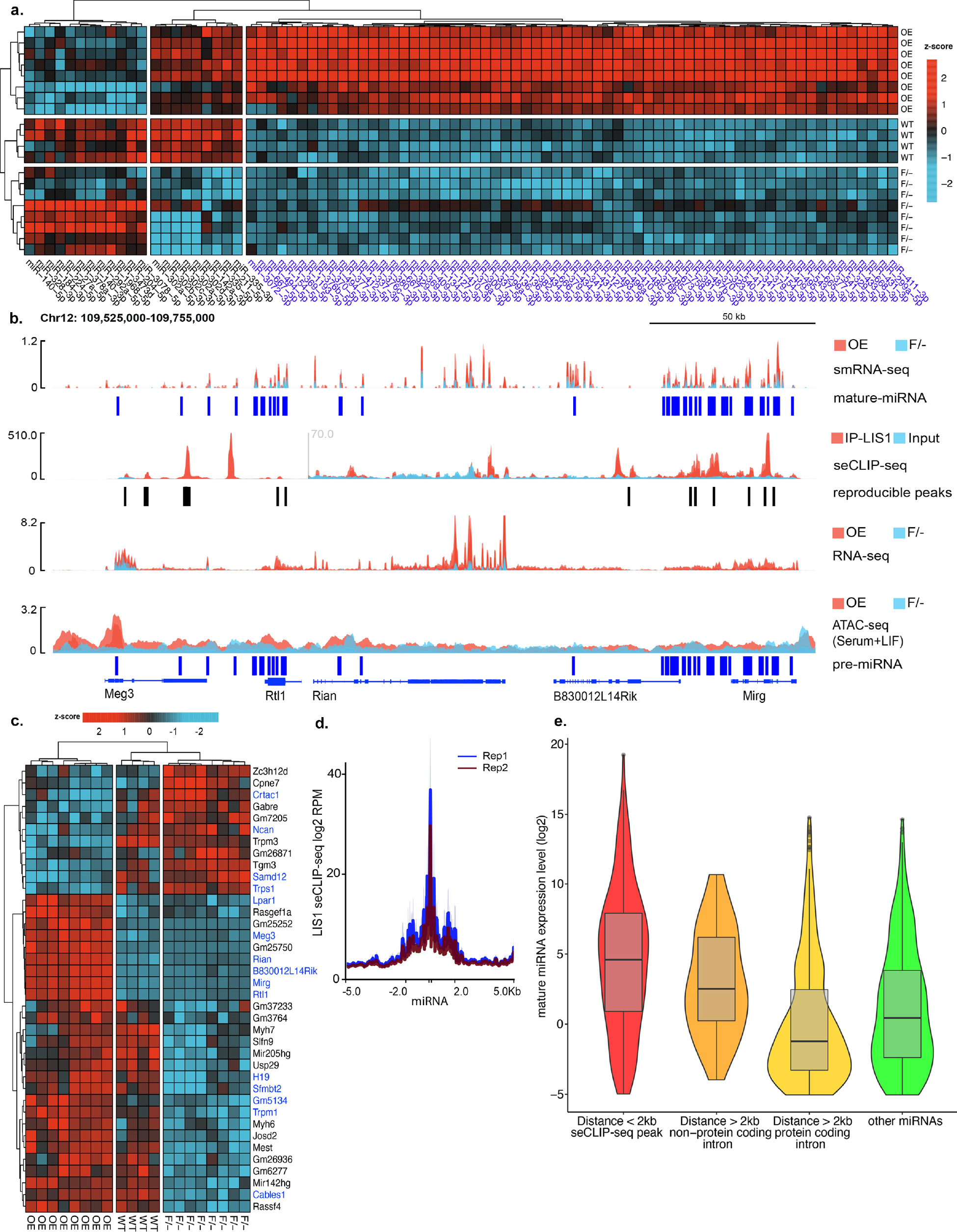
I**n**creased **LIS1 expression drives the expression of miRs. a.** A heatmap of the top 85 DE mature miRs in comparing LIS1 OE and F/- mESCs with WT is shown. The data are shown on a Z-score scale of the variance stabilizing transformation on normalized reads. **b.** *Meg3-Mirg* locus. Top to bottom tracks: small RNA-seq signal (OE in red and F/- in blue); seCLIP-seq (IP in red and input in blue); bulk RNA-seq (OE in red and F/- in blue); ATAC-seq (OE in red and F/- in blue). All tracks are normalized. Of note, the Meg3 promoter region shows significant differential accessibility between the OE and F/- samples. The plot shows the merged track for replicates of each mESCs non-isogenic clone. **c.** A heatmap of differentially expressed miR host genes from bulk RNA-seq. The data are shown on a Z-score scale of the variance stabilizing transformation on normalized reads. **d.** Metagene plot of LIS1 seCLIP-seq coverage as a function of distance from all pre-miRs in the mouse genome. **e.** Expression (log2 DESeq2 baseMean) of miRs as a function of their distance from the closest LIS1 seCLIP-seq peak. Red; LIS1 seCLIP-seq peak in a distance < 2000bp, Orange; LIS1 seCLIP-seq peak with a distance > 2000bp, the peak resides in intron of non-coding gene, Yellow; LIS1 seCLIP-seq peak with a distance > 2000bp, the peak resides in intron of protein-coding gene, Green; all other miRs.

**Fig. 5:**
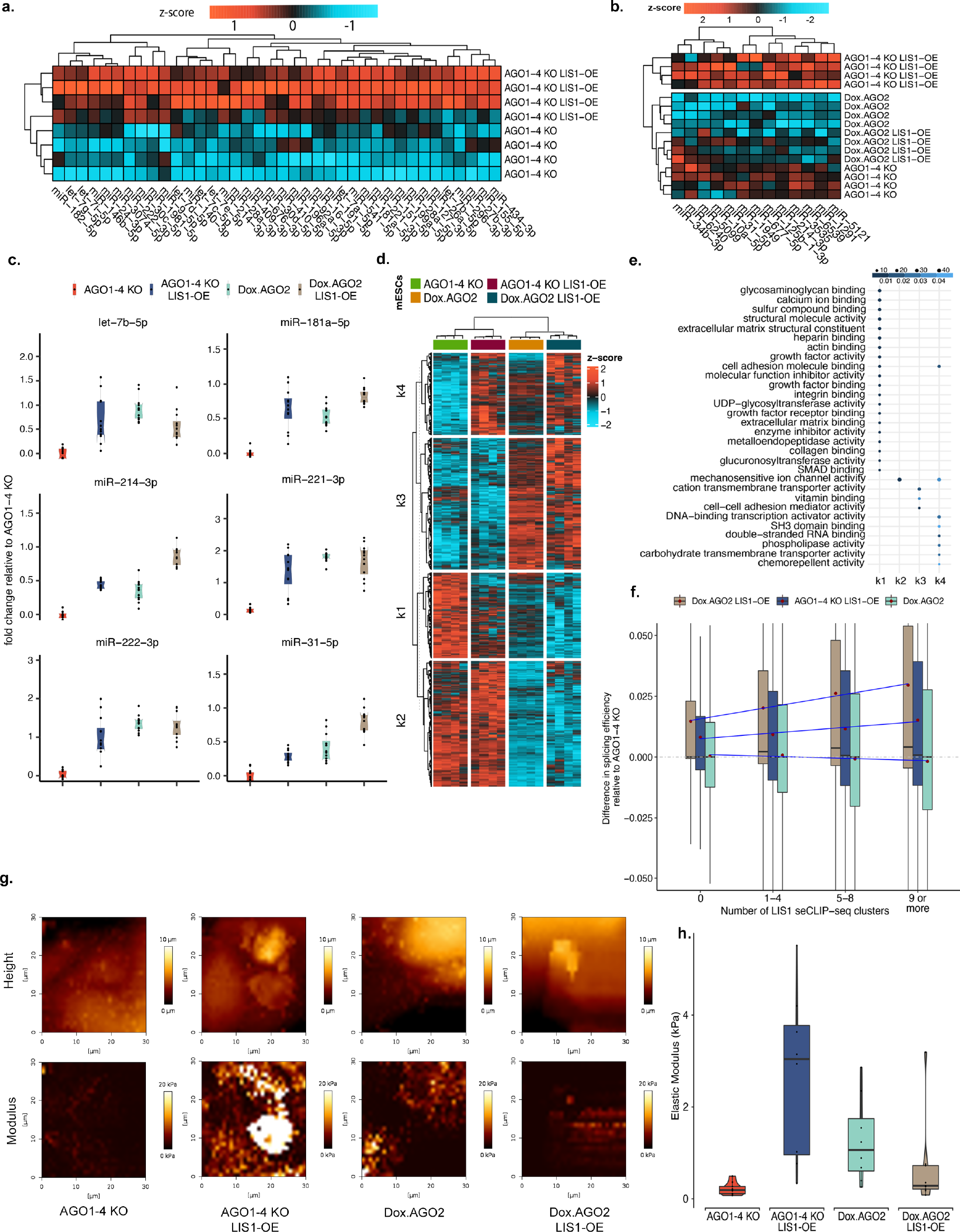
L**I**S1 **OE affects gene expression and cell stiffness in AGO1-4 KO mESCs. a.** A heatmap showing the increase in mature miRs’ expression due to LIS1 overexpression (LIS1 OE ^AGO1-4KO^) compared to AGO1-4 KO. The scale bar represents the z-score (p ≤ 0.05, fold change ≥ 0.5). **b.** A heatmap showing the increase in mature miR expression due to LIS1 overexpression (LIS1 OE ^AGO1-4KO^) compared to Dox. AGO2 ^AGO1-4KO^ with AGO1-4KO and Dox. AGO2 LIS1- OE ^AGO1-4KO^. For **a.** and **b.,** the data are shown on a Z-score scale of the variance stabilizing transformation on normalized reads. **c.** qRT-PCR for a subset of mature miRs involved in regulating ECM and mechanosensitive genes (n = 4 (x3), p-values are reported in extended data table 6). **d.** A heatmap of 3183 genes differentially expressed due to LIS1 overexpression and doxycycline induced AGO2 expression in AGO1-4 KO mESCs cultures. The data are shown on a Z-score scale of the variance stabilizing transformation on normalized reads. **e.** Analysis of gene set over-representation test for four clusters obtained from the k-means clustering, illustrating the combined effect of LIS1 and AGO2 modulating the expression of genes in top enriched Gene Ontology Molecular Function (GO-MF). **f.** Intron level quantification of differences in splicing efficiency for Dox. AGO2 ^AGO1-4KO^, LIS1-OE ^AGO1-4KO,^ and Dox. AGO2 LIS1-OE ^AGO1-4KO^ relative to AGO1-4KO and correlated to the number of LIS1 seCLIP-seq clusters. All the comparisons were significant across each genotype in each cluster group, and p- values for significance are reported in extended data Table 6. **g.** Atomic force microscopy images show on the top height and the bottom the elastic modulus of stem cell colonies. Left to right- AGO1-4KO, Dox. AGO2 ^AGO1-4KO^, LIS1-OE ^AGO1-4KO,^ and Dox. AGO2 LIS1-OE ^AGO1-4KO^, respectively (scale bars, five μm). **h.** Box plots showing Young’s modulus (in kilopascals (kPa), y-axis) for independent and combined effects of constitutive LIS1-OE and Dox.AGO2 in AGO1- 4 KO mESCs. Boxes show the median and distribution of median absolute deviation values for the measurements of each group. A histogram of modulus values for each measurement (6-8 colonies per group) is shown in extended data Fig. 9.

### LIS1 affects the expression of miRs

We next proceeded to examine the effect of LIS1 on small RNA. Argonaute proteins are best-known for their role in post-transcriptional regulation by microRNAs (miRs)^57^; therefore, we conducted RNA-seq and small-RNA seq from mESCs. We detected 85 DE mature miRs, most of which were upregulated by LIS1 overexpression (Fig. 4a, Extended Data Fig. 6a,b, Extended Data Table 10). Among the DE miRs, the *Meg3-Mirg* locus was highly represented and included 76 small RNA genes (64 mature miRs) (Fig. 4b). Multiple LIS1 seCLIP peaks were detected in this locus, many of which were on top of or in close proximity to miRs (Fig. 4B). Not only was the expression of miRs in this locus significantly increased but also the expression of the protein-coding and non-protein-coding genes (Fig. 4c, Bulk RNA-Seq data in Extended Data Table 11). ATAC-seq experiments revealed that this locus and the associated enhancer region are relatively open in the LIS1 OE, facilitating transcription (Fig. 4b). Most of the DE miRs were located in introns or in close proximity to protein-coding or non-coding genes, many of which were also DE (Fig. 4c). The expression of the majority of these genes was elevated in the LIS1 OE line (Fig. 4c). We further correlated the expression of miRs with known mRNA targets in our RNA-seq data (Extended Data Fig. 6c). We noted that the expression of miRs in close proximity to LIS1 seCLIP peaks is significantly higher than those not located in proximity to the LIS1 seCLIP peaks (Fig. 4d,e). We then examined the expression of miRs in relation to LIS1 seCLIP peaks using a threshold of 2kb (Fig. 4e, statistics in Extended data Table 7). The expression of a miR located in the LIS1 seCLIP site (±2kb) was significantly higher than in all other categories. If the location of the miR was further away from the peak (>2kb), their level of expression was dependent on whether they were in non-coding or protein-coding introns, with those located in introns of non-coding RNAs having higher expression than those found in protein-coding introns or in comparison to all other miRs (statistics in Extended Data Table 7).

Collectively, our data indicate that increased levels of LIS1 stimulate the expression of a subset of miRs. LIS1’s binding to intronic sequences on top of or near miRs is correlated with higher expression of these miRs. Still, if the distance to the LIS1 seCLIP binding site is beyond 2 kb, the miRs located in introns of non-protein-coding genes will tend to be expressed. In contrast, those found in introns of protein-coding genes are likely not to be expressed in the mESCs. LIS1 likely affects the expression of miRs at multiple levels ^58^.

### LIS1 affects the expression of miRs without the Argonaute complex

To further interrogate the interactions between LIS1/AGO2 and the microRNA pathway, we generated an additional cell line where LIS1 is overexpressed (LIS1 OE) in the context of cells lacking Argonaute proteins (LIS1 OE AGO1-4 KO) with the possibility to induce AGO2 expression (LIS1 OE Dox AGO2). The AGO1-4 KO line exhibited a paucity of miRs as previously described ^53^, yet LIS1 OE was sufficient to slightly but significantly increase the expression of a subset of miRs (Fig. 5a, Extended Data Table 12). An additional small subset of miRs whose expression was slightly but significantly increased following LIS1 OE was noted in the background of AGO2 expressing cells (Fig. 5b). The relative expression of a subset of miRs was examined by qPCR (Fig. 5c, Extended Data Fig. 7a, statistics in Extended Data Table 7).

Among the 85 DE miRs we previously identified, many are known to regulate tissue stiffness and be involved in mechanosensitivity ^59–63^, including several members of the let-7 family, miRs 221/222-3p, 146-5p, and 151-5p. In addition, we noted the increased expression of all of the members of the miR 302 family, known to be involved in the pluripotency network in mESCs ^64^.

The expression of these miRs and others significantly increased in the presence of LIS1 OE (Fig. 5c, Extended Data Fig. 7a). The expression of the let-7 family of miRs may be associated with the fact that they are negatively regulated by LIN28a ^65^, and this protein is part of the LIS1 interactome (Fig. 2 and Extended Data Table 5). As indicated above, LIS1 plays a role in splicing. Therefore, we examined the differences in splicing efficiency of miR host genes at the transcript and intron levels in all the cell lines compared to the AGO 1-4KO line (Extended Data Fig. 7 b-c). Significant changes were noted in all comparisons, yet the p values were more pronounced when the intron levels were examined. The heatmap of bulk RNA-seq depicted that the expression of AGO2 introduced a significant change to gene expression (Fig. 5d, Extended Data Table 13). However, in the LIS1 OE AGO1-4 KO, we noticed that the expression pattern of a subset of the genes was similar to that in the DoxAGO2, suggesting that LIS1 OE can partially rescue the dysregulation in AGO1-4 KO cells (Fig. 5d). The GO molecular function terms of the DE genes indicated multiple ECM functions and mechanosensitivity (Fig. 5e).

We then examined local splicing variations (LSV) by comparing the different AGO lines with and without LIS1 OE using MAJIQ (Extended Data Fig. 8a,b, Extended Data Table 14).

Reactome pathway enrichment analysis for differentially spliced genes revealed signaling by TGF, RHO GTPase cycle, and chromatin organization (Extended Data Fig. 8c). Examples for alternative splicing events are shown for *Meg3* and *Rian* (Extended Data Fig. 8d,e). We then examined the differences in splicing efficiency between AGO1-4 KO and the rest of the lines in relation to the number of LIS1 seCLIP clusters (Fig. 5f, statistics in Extended Data Table 7). In the case of the two lines that overexpressed LIS1, we noted that the differences in splicing efficiency increased with the gain in LIS1 seCLIP clusters. The difference in splicing efficiency following induction of AGO2 expression in the AGO1-4 KO was not affected by the presence of LIS1 RNA-binding clusters.

The RNAseq and small RNAseq suggested that the different mESC lines on the AGO1-4 KO background likely differ in their physical properties. To test this hypothesis, we subjected mESC colonies to Atomic Force Microscopy (AFM) nanomechanical measurements (Fig. 5g-i, Extended Data Fig. 9, statistics for 5h is in Extended Data Table 7). The elastic modulus of AGO1-4 KO was the lowest; the addition of LIS1 to these cells (AGO1-4 KO LIS1 OE) exhibited the highest value. The changes in the stiffness are correlated with changes in gene expression that were observed above (Fig. 5d-e). The Dox AGO2 value was higher than that of the AGO1-4 KO.

Collectively, our data indicate that LIS1 OE can increase the expression of a subset of miRs and can affect gene expression and splicing in an AGO2 independent way. Furthermore, LIS1 OE, together with AGO2 or independent of AGO2, can modulate the stiffness of mESC.

## Discussion

LIS1 has been studied intensively for several decades ^66^, yet here we show a myriad of novel roles for this protein, especially in relation to post-transcriptional regulation. RNA-seq from LIS1 null cells, cultured in a custom-designed media, indicated the involvement of LIS1 in the regulation of gene expression related to many dynein-mediated activities such as mitosis and microtubule-based transport, adding another level of regulation of dynein functions by LIS1. In addition, we also detected pathways related to organophosphate biosynthetic and fatty acid metabolic processes that were captured in our metabolomics analysis. Several of the affected pathways were related to RNA. LIS1 dosage also affected gene expression in hESC, with a striking absence of E-cadherin in *LIS1-/+* hESCs evident by immunostainings. Loss of E-cadherin can promote epithelial to mesenchymal transition, affecting the stem cells’ pluripotency ^67^. To further understand the molecular mechanisms involved in these diverse activities, we have undertaken an unbiased approach to compile the LIS1 interactome in the nuclei and cytoplasm of mESCs with different levels of LIS1. The known LIS1 interactome included only 148 proteins that are mainly involved in regulating the cytoskeleton. Our experiment revealed that the LIS1 interactome is composed of several hundred proteins. Many LIS1-interacting proteins are known RBPs, with a striking representation of the Argonaute complex. We further dissected the LIS1 interactome to AGO2-dependent and -independent subgroups. The enrichment of RBPs in the LIS1 interactome combined with a previous high throughput study that identified LIS1 as an RNA-binding protein ^54^ warranted further analysis using seCLIP. LIS1 was bound mainly to introns of nascent RNA of protein-coding genes. The presence of an increased number of binding sites within a gene was negatively correlated with its’ splicing efficiency, while an increased level of LIS1 was positively correlated with splicing efficiency. The same trend was noted when LIS1 was overexpressed in the background of AGO1-4 KO and Dox AGO2 cell lines. We noted that changes in LIS1 dosage affected splicing in multiple genes, resulting in skipped exons or retained introns, which can have various effects on steady-state RNA levels and protein translation ^68^.

The association of LIS1 with the Argonaute complex strongly suggests that the expression of miRs might be affected by LIS1 dosage. Furthermore, the LIS1 interactome shares fifty-seven proteins in common with a list of 181 proteins detected in a large-scale biochemical screen to identify a comprehensive list of RBP-miRNA interactions ^69^ (significant overlap using a hypergeometric test; representation factor: 2.8, p < 1.980e-13). These RBPs are highly enriched for proteins involved in RNA splicing. Therefore, the effect of LIS1 could be mediated through the RNA-induced silencing complex (RISC) or other LIS1 interacting RBPs. In the context of LIS1 dosage, LIS1 OE significantly increased the expression of multiple miRs, many of which were located in operons. Suboptimal Drosha/DGCR8 substrate miRNAs are enriched in operons, and their proximity facilitates their subsequent processing ^70^. The protein-coding and non-protein-coding genes in the DE miR loci were also DE. This correlation may be related to the observed changes in chromatin organization evident by ATAC-seq. The expression level of miRs was positively correlated with their distance from LIS1 seCLIP sites. Therefore, our data strongly suggest that LIS1 is involved in positive regulation of the expression of these miRs either by recruitment of additional proteins to the nascent RNA or by affecting RNA splicing that is involved in the formation of those miRs ^71^. However, other mechanisms are likely to be engaged in DE miRs. It has been proposed that impaired export of miRs via exosomes may increase cellular miRs ^72^. Considering the known roles of LIS1 in intracellular transport, LIS1 may affect miRs at multiple levels. Most studies focus on the stages of miR biogenesis, but only a few studies investigated the RISC stage. The RISC loading involves the binding of Argonautes to miRs or siRNA duplexes. The passenger stands of the miRs or the siRNA duplexes are degraded, and the mature RISC is guided to the respective mRNA, or Argonautes binds to preformed duplexes of miR/siRNA - mRNA ^73, 74^. Several proteins are involved in the formation of these AGO-independent duplexes. For example, the RBP AUF1 (HNRNPD) can directly promote the binding of the miRNA let-7b to its target site within the 3ʹUTR of the POLR2D mRNA ^75^. In a systematic eCLIP study examining a set of 126 RBPs, the vast majority (92%) interacted with at least one miR locus^76^, suggesting that our understanding of how RBPs regulate miR expression is far from complete.

Our findings indicated that LIS1 OE could increase the stiffness of Argonaute deficient cells. It has been previously demonstrated that AGO2 activity is required for tissue stiffness ^59^, but it was unknown if the physical properties of the cells can change in an AGO2 independent manner.

LIS1 OE in the absence of AGO1-4 resulted in a slight but significant increase in a small set of miRs. Data mining revealed that many of these miRs, including members of the let-7 family, miR-221-3p, 16-5p, 296-3p, and others, are involved in mechanotransduction pathways ^60, 77–85^. RNA seq data further revealed the DE of genes associated with mechanosensitive pathways.

Indeed, in the absence of all Argonaute proteins, LIS1 OE induces a significant elevation in the measured Young’s modulus.

Taken together, our studies have demonstrated that changes in LIS1 expression modulate chromatin organization, RNA and small RNA gene expression, and splicing, which results in long-lasting changes in the physical properties of mESCs. We propose that these previously unrecognized, nuclear, RNA- and small-RNA-related LIS1 activities underlie, at least in part, some of the activities that make LIS1 so crucial for proper human brain development ^1, 3, 66^.

## Acknowledgments

We thank Samara Brown, current and previous members of the Reiner and Hanna lab. Merav Kedmi and Muriel Chemla from the Biological services; Current and ex-staff from the INCPM for help with the library prep and sequencing. Yungui Yang for help with sequencing at the Beijing genomics institute. Eran Hornstein and Eli Arama for helpful discussions. Phillip Sharp for the AGO mESCs.

## Author contributions

Conceptualization: AK, OR, TS

Methodology: AK, TO, AG, TS, DT, IRG, SM, MI, AA, YMG, SRC, IU

Visualization: AK, TO, IU, AA, IRG, SM, SI Funding acquisition: OR, TS, KK

Project administration: TO, TS, JH, YMG, KK, OR Supervision: OR, TO, IU, KK

Writing – original draft: AK, OR, TO, TS

Writing – review & editing: AK, AG, TO, TS, DT, IRG, SM, MI, AA, YMG, SRC, JH, IU, KK, OR

## Data and materials availability

All data, code, and materials used in the analysis is available per request. Stem cell lines and plasmids generated in this study require materials transfer agreements (MTAs). All the omics studies are deposited in public databases, and the accession numbers are pending. Most of the data are available in the main text or supplementary materials.

## Methods

### Generation of mESC lines

All animal studies were done following the approval of the IACUC committee at the Weizmann Institute of Science. The use of experimental animals is in complete accordance with: The Animal Welfare Law (Experiments with animals); The Regulations of the Council for Experiments with Animals; The Weizmann Institute Regulations (SOP); The Guide for the Care and Use of Lab Animals, National Research Council, 8th edition; The Guidelines for the Care and Use of Mammals in Neuroscience and Behavioral Research.

To generate LIS1 mutant mESCs, the *Lis1* F/F (*Lis1^flox/flox^*) mice were crossed with mice expressing tamoxifen-inducible Cre with a ubiquitin promoter (*UB-Cre/ERT2*) or with a constitutive maternal Cre expressed under the regulation of the phosphoglycerate kinase promoter (PGK-Cre). *Lis1* +/- without PGK-Cre were crossed with *UB-Cre/ERT2*:*Lis1* F/F to get *UB-Cre/ERT2*:*Lis1* F/- and *Lis1* F/-. LIS1-FLAG-DsRed mice^3^ were crossed with PGK-Cre mice to obtain PGK Cre: *Lis1*+/+: LIS1-FLAG-DsRed to obtain ubiquitous LIS1 transgene overexpression. Blastocysts were flushed at embryonic day 3.5 (E3.5) in KSOM medium (Invitrogen, Thermo Fisher Scientific) from naturally mated timed-pregnant mice and cultured for five days in 2i+LIF medium (Supplementary Table1). All mESCs were routinely maintained on irradiated MEFs in FBS+LIF medium (Supplementary Table1). To ensure stable LIS1 overexpression in *UB-Cre/ERT2*:*Lis1* F/- and TT-FHAgo2 ^87^ (received from Drs. Zamudio, Suzuki, and Sharp), the PiggyBac transposase system was used^88^. To generate *UB- Cre/ERT2*:*Lis1-F/-:*LIS1-GFP-OE [LIS1-GFP-OE (F/-)] and TT-FHAgo2:LIS1-GFP-OE

(AGO1-4 KO LIS1-OE), we transfected 10μg of a mix containing the pCAG:*LIS1*-GFP plasmid with a pCAG-PBase plasmid expressing the PiggyBac transposase. The transfection was done using a NEPA21 electroporation system according to the manufacturer’s instructions. Following electroporation, cells were passaged once after five days, and GFP-positive cells were selected using fluorescence-activated cell sorting (FACS). To induce the deletion of *Lis1*, cells were treated with 500nM 4-hydroxytamoxifen (4-OHT) for 72h in 5i+LIF (Supplementary Table1) with 100 μM apoptosis inhibitor Z-vad (Sigma-Aldrich®) and 50 μM of necroptosis inhibitor Necrostatin-1 (Sigma-Aldrich®), respectively. The deletion was assessed by genotyping for the floxed and deleted alleles. For mESCs grown in 5i+LIF, the medium was replaced daily. TT- FHAgo2 and TT-FHAgo2 LIS1-GFP-OE clonal cell lines were maintained in FBS+LIF medium and were treated for 72hr with 0.1 μg/ml doxycycline to induce AGO2 expression. We used the naive pluripotency reporter mESCs line V6.5 deltaPE-Oct4-GFP ^89–91^ which were grown in 5i+LIF (3-4 passages), colony morphology was assessed compared with cells cultured in other media (Supplementary Table 1).

WIBR3 (NIHhESC-10-0079) hESCs and isogenic *LIS1*+/- ^10^ were grown and maintained on irradiated MEFs in optimal naive NHSM conditions (RSET-Stem Cell Technologies INC, supplementary Table 1) ^90^. Gain of function LIS1-GFP-OE isogenic hESCs were generated with the PiggyBac pCAG:*LIS1*-GFP transfection in WIBR3 hESCs as described above. LIS1 int6/int6 homozygous mutation isogenic hESCs were generated using the CRISPR/Cas9 protocols previously described, with some modifications. Briefly, to introduce cis-splicing mutation, the CRISPR genome-editing method with double Cas9 nickase was used to reduce the probability of off-target effects ^92^. The two guide-RNA sequences, sense: ATATTGCTGTTATGTGTTTT and antisense: TGGCTACTGAAGAAACATTG, were designed according to http://crispr.mit.edu/ and were cloned into a pX335 vector ^92^ targeting intron 6 near the acceptor site of exon 7 of the *LIS1* gene. Cells were transfected using electroporation as described above with a repair single-strand oligo cccatggtcaattgatgtttcattgctcttggtggtatattacttcataatatattgctgttaCgtgttttagGCCATGAtCACAATGTTTCTTCAGTAGCCATCATGCCCAATGGAGATCATATAGTGTCTGCCTCAAGGGATAA

AAC encoding the T>C mutation and introducing a silent mutation leading to a new BclI site and the pX335 plasmid with trace amounts of a GFP expression vector. Three days after transfection, the cells were subjected to FACS and plated at a density of 2,000 cells per 10 cm plate on irradiated MEFs, allowing for the growth of single-cell-derived colonies. Clone 61 was used in this study. The mutation was confirmed by PCR, restriction enzyme digestion, and Sanger DNA sequencing. The hESCs were passaged with TrypLE (Thermo Fisher Scientific) every 3-4 days and were grown on irradiated MEFs for four passages in tHENSM ^35^ when the media was changed from NHSM (supplementary Table 1). *LIS1*+/- hESCs lose colony morphology and do not grow well after five passages in tHENSM media.

### Pre-implantation embryo and cell lines immunostaining

E3.5 WT embryos were collected in KSOM media, and after three washes with PBS, embryos were fixed with 4% PFA overnight at 4°C. The staining procedure was performed in Pyrex spot plates under a Delta vision microscope. ESCs were cultured in multiwell glass-bottom plates (MatTek Life Sciences) coated with Matrigel (Corning Life Sciences) and were fixed with 4% PFA for 20 minutes at room temperature. Samples were rinsed thrice in PBS and were permeabilized in permeabilization solution (0.1% Triton X-100 in PBS) for 20 minutes at room temperature. This was followed by incubation in blocking solution (2% donkey serum or 2% normal goat serum, 0.1% BSA in permeabilization solution). Samples were incubated overnight with primary antibodies in blocking solution (1:100). On the next day, samples were rinsed three times with blocking solution for 10 minutes each. This was followed by incubation with secondary antibodies diluted in blocking solution (1:500) for 1 hour at room temperature and costained with DAPI (1 μg/ml in PBS) for 5 minutes. Embryos were moved and allowed to sink in 96 well MatTek plates in PBS. Imaging was done using a spinning disk confocal microscope based on an OLYMPUS IX83 inverted microscope, VisiScope CSU-W1-T1 confocal system (Visitron Systems, Germany), and an sCMOS 4.2 MPixel camera. Imaging was performed using the VisiView software. Imaging of ESCs was carried out using a Dragonfly 200 spinning disk confocal microscope (ANDOR, Oxford instruments). Images were processed using Imaris microscopy image analysis software (Oxford instruments). The following antibodies were used: anti-LIS1 (Sapir T. et al.^26^, 338), anti-Nanog (AF2729, R&D), anti-OCT3/4 (C10, Santa Cruz), anti-E-CADHERIN (Abcam, Cambridge, UK), and anti-Pan-LEFTY (Abcam, Cambridge, UK).

### Metabolite extraction

mESCs were grown in FBS+LIF medium without Dimethyl 2-oxoglutarate. Cells were trypsinized and centrifuged at 500*g* for 3 min. The medium was completely removed, and 6x10^6^ feeder MEFs depleted ESCs were taken for sample preparation. Extraction and analysis of lipids and polar metabolites was performed as previously described ^93, 94^ with some modifications: samples were mixed with 1 ml of a pre-cooled (−20°C) homogenous methanol:methyl-tert-butyl-ether (MTBE) 1:3 (v/v) mixture, containing following internal standards: 0.1 μg/ml of Phosphatidylcholine (17:0/17:0) (Avanti), 0.4 μg/ml of Phosphatidylethanolamine (17:0/17:0, 0.15 nmol/ml of Ceramide/Sphingoid Internal Standard Mixture I (Avanti, LM6005), 0.0267 µg/ml d5-TG Internal Standard Mixture I (Avanti, LM6000) and 0.1 μg/ml Palmitic acid-13C (Sigma, 605573). The tubes were vortexed and then sonicated for 30 min in an ice-cold sonication bath (taken for a brief vortex every 10 min). Then, double deionized water (DDW): methanol (3:1, v/v) solution (0.5 ml) containing the following internal standards: C13 and N15 labeled amino acids standard mix (Sigma) was added to the tubes followed by centrifugation. The upper organic phase was transferred into a 2 ml Eppendorf tube. The polar phase was re-extracted as described above, with 0.5 ml of MTBE. Both parts of the organic phase were combined and dried in speedvac and then stored at −80°C until analysis. For analysis, the dried lipid extracts were resuspended in 150 μl mobile phase B (see below) and centrifuged again at 13,000 rpm at 4°C for 5 min. The lower polar phase was lyophilized and stored at −80°C until analysis. Before the injection, the polar phase samples pellets were dissolved using 150 ul DDW- methanol (1:1), centrifuged twice (13,000 rpm) to remove possible precipitants, transferred to HPLC vial, and were injected into the LC-MS system.

### LC-MS for lipidomics analysis

Lipid extracts were analyzed using a Waters ACQUITY UPLC system coupled to a Vion IMS qTof mass spectrometer (Waters Corp., MA, USA). Chromatographic conditions were as described^93^ with small alterations. Briefly, the chromatographic separation was performed on an ACQUITY UPLC BEH C8 column (2.1×100 mm, i.d., 1.7 μm) (Waters Corp., MA, USA). The mobile phase A consisted of DDW: Acetonitrile: Isopropanol 46:38:16 (v/v/v) with 1% 1 M NH4Ac, 0.1% glacial acetic acid. Mobile phase B composition is DDW: Acetonitrile: Isopropanol 1:69:30 (v/v/v) with 1% 1 M NH4Ac, 0.1% glacial acetic acid. The column was maintained at 40°C; the flow rate of the mobile phase was 0.4 ml/min, the run time was 25 min. The linear gradient was as follows: Mobile phase A was run for 1 min at 100%, then reduced to 25% for 11 min, followed by a decrease to 0% for 4 min. Then, mobile phase B was run at 100% for 5.5 min, followed by setting mobile phase A to 100% for 0.5 min. Finally, the column was equilibrated at 100% A for 3 min. MS parameters were as follows: the source and de-solvation temperatures were maintained at 120 °C and 450 °C, respectively. The capillary voltage was set to 3.0 kV and 2 kV for positive and negative ionization mode, respectively; cone voltage was set for 40 V. Nitrogen was used as de-solvation gas and cone gas at a flow rate of 800 L/h and 30 L/h, respectively. The mass spectrometer was operated in full scan HDMS^E^ resolution mode over 50–2000 Da mass range. For the high-energy scan function, a collision energy ramp of 20–80 eV was applied; for the low energy scan function, 4 eV was applied.

### Lipid identification and quantification

LC-MS data were analyzed and processed with UNIFI (Version 1.9.3, Waters Corp., MA, USA). The putative annotation of the lipid species was performed by comparison of accurate mass (below 5 ppm), fragmentation pattern, retention time (RT), and ion mobility (CCS) values to an in-house-generated lipid database. Peak intensities of the identified lipids were normalized to the internal standards and the amount of protein in the cells, used for analysis.

### LC-MS polar metabolite analysis

Metabolic profiling of the polar phase was described^11^ with minor modifications described below. Briefly, analysis was performed using Acquity I class UPLC System combined with mass spectrometer Q Exactive Plus Orbitrap™ (Thermo Fisher Scientific), which was operated in a negative ionization mode. The LC separation was done using the SeQuant Zic-pHilic (150 mm × 2.1 mm) with the SeQuant guard column (20 mm × 2.1 mm) (Merck). The Mobile phase B: acetonitrile and Mobile phase A: 20 mM ammonium carbonate with 0.1% ammonia hydroxide in DDW: acetonitrile (80:20, v/v). The flow rate was kept at 200 μl/ min and gradient as follows: 0-2min 75% of B, 14 min 25% of B, 18 min 25% of B, 19 min 75% of B, for 4 min.

### Polar metabolites data analysis

Data processing was done using TraceFinder (Thermo Fisher Scientific) when detected compounds were identified by accurate mass, retention time, isotope pattern, fragments and verified using an in-house-generated mass spectra library.

### Immunoprecipitation and mass spectrometry

To identify LIS1 interacting proteins from wild type (WT), Floxed/- (F/-), and LIS1- dsRED OE (OE) mESC, we prepared cytoplasmic and nuclear extracts. mESCs cultured in 15cm culture dishes were washed with ice-cold PBS. Cells were gently scraped on ice. The cytoplasmic fraction was extracted by incubating in Buffer A (10 mM HEPES pH 7.8, 10 mM KCl, 2 mM MgCl2, 10 mM NaF, 0.05% NP40) supplemented with a protease inhibitor cocktail (Sigma). After centrifugation (for 5 min at 500 *g* at 4°C), the pellet (the nuclear fraction) was resuspended in twice the volume of high salt buffer (20 mM HEPES pH 7.8, 0.6 M KCl, 2 mM MgCl2, 25% glycerol, 10 mM NaF, protease inhibitor cocktail), incubated on ice for 30min, and centrifuged for 30min at 24000 g at 4°C. For immunoprecipitation (IP), Eppendorf tubes containing A/G protein beads were incubated with monoclonal anti-LIS1 antibodies (20 μl for each sample, 338) in blocking buffer (1xPBS, 0.5% Tween-20, 0.5% BSA) and rotated for 1h at 25°C. Next, the nuclear lysate was added and incubated for 6 hours at 4°C in IP buffer (50 mM Tris-HCl pH 7.5, 150 mM NaCl, 1% Triton X-100 and protease inhibitor cocktail) while rotating. Immunoprecipitated proteins were pelleted by centrifugation and washed three times with IP buffer. To identify LIS1 interacting proteins from AGO1-4 KO and Dox.AGO2 mESCs whole-cell lysates (cytoplasm and nuclear extracts) were prepared. Lysates were immunoprecipitated with anti-LIS1 antibodies or IgG antibodies as described above.

Mass spectrometry was performed as previously described^95^ with some modifications. Following the IP with anti-LIS1 antibodies, the precipitated proteins were extracted from the beads using an SDS-sampling buffer. We employed a FASP method to remove SDS from the solubilized protein^96^. Briefly, the SDS samples were applied onto an Amicon filter unit (MWCO = 10 kDa; Ultracel-10 cellulose membrane, Cat.No; UFC201024, Merck Millipore, Billerica, MA, USA) to trap the proteins. After washing the filter units with 8-fold volume (v/v) of Urea buffer (100 mM Tris-Cl (pH8.5), 8 M Urea), the trapped proteins on the filter membrane were alkylated using 10 mM iodoacetamide for 1hr in the dark. The proteins were digested with Trypsin/ Lys-C (Promega, WI, USA). According to the manufacturer’s instructions, demineralization was performed using SPE c-tips (Nikkyo Technos, Tokyo, Japan). The peptides were analyzed by LC/MS using an Orbitrap Fusion mass spectrometer (Thermo Fisher Scientific Inc, MA, USA) coupled to an UltiMate3000 RSLCnano LC system (Dionex Co., Amsterdam, the Netherlands) using a nano HPLC capillary column (Nikkyo Technos Co., Tokyo, Japan) via a nanoelectrospray ion source. Reversed-phase chromatography was performed with a linear gradient (0 min, 5% B; 100 min, 40% B) of solvent A (2% acetonitrile with 0.1% formic acid) and solvent B (95% acetonitrile with 0.1% formic acid) at an estimated flow rate of 300 nl/min.

### Mass spectrometry data analysis

Mass spectrometry raw files were processed using MaxQuant software (version 1.6.2.10)^97^ and the Andromeda search engine. Database searches were done against the reference proteome of Mus musculus obtained from UniProtKB in November 2016. The carbamidomethylation of cysteine was set as a fixed modification, and the oxidation of methionine, the phosphorylation of Ser/Thr/Tyr, and N-terminal acetylation were set as variable modifications. Trypsin without proline restriction enzyme option was used, with two allowed miscleavages. False discovery rates (FDRs) for the peptide, protein and site levels were set to 0.01 with minimal unique+razor peptides number set to 1. The sum of unique+razor peptides across replicates from any of the fractions with the minimum of two peptides higher than control were considered candidate substrates. Peptide intensity normalization and quantification were performed with the limma algorithm in the DEP Bioconductor package (v.1.16.0)^98^. Mixed imputation was performed on proteins by using a linear model with MAR = zero and MNAR = MinProb. Pairwise differential testing was performed using test_diff function, and significant proteins were identified using add_rejection function, setting p-value threshold (alpha) 0.05. For the ortholog similarity, *Drosophila melanogaster* and *Mus musculus* ensemble gene pair list was obtained for FlyBase gene identifiers. These were matched with *Mus musculus* homolog-associated gene name using Bioconductor package biomaRt (v2.50.1).

### RNA-seq library preparation

Total RNA was extracted using the miRNeasy Mini kit (Qiagen, Hilden, Germany), and any residual genomic DNA was removed using a DNA-free DNA Removal Kit (Qiagen, Hilden, Germany). RNA concentration and integrity were measured using Nanodrop (Thermo Scientific) and an Agilent Tapestation 4200, respectively. Total RNAseq libraries from 1μg of ribosomal depleted RNA using TruSeq Stranded Total RNA Library Prep (Illumina) or Lexogen SENSE mRNA-Seq Library Prep Kit. These libraries were sequenced on Illumina NextSeq 550 or HiSeq X-Ten to obtain 125bp or 150bp paired-end libraries. As previously described, bulk MARS-seq libraries were produced from a modified version of 3’ end single-cell RNA-seq^99^. Sequencing libraries were prepared from 50ng of purified total RNA and were pooled for biological replicates. The quality of the final libraries was assessed with qPCR and Agilent TapeStation and was sequenced in the Illumina NextSeq 550 sequencer to obtain a single read of 75bp.

### RNA-seq analysis

RNA-seq reads were processed using the UTAP pipeline^100^. Briefly, fastq files trimmed with cutadapt and aligned to the GRCm38/mm10 reference genome using STAR (version v2.4.2a), with parameters –alignEndsType EndToEnd, outFilterMismatchNoverLmax 0.05, – twopassMode Basic. Gene read count was performed using the qCount function from the QuasR Bioconductor package (v.1.34)^101^ with default parameters. The batch effect of the RNAseq read count was corrected using Combat-seq^102^ with a negative binomial regression model on the raw read count data, from the sva Bioconductor package (v.3.42). The corrected read count was then used for testing differential expression with DESeq2^103^ Bioconductor package (v.1.34).

Estimation of group size differences was performed with lfcShrink function with -type apeglm^104^. For MARS-seq, samples were analyzed using the UTAP pipeline^100^. Reads were trimmed and aligned to the GRCm38/mm10 and GRCh38/hg38 reference genome for mouse and human, respectively, as described above. Gene quantification of the most 3’ 1000bp of each gene was performed using HTSeq-count^105^ in union mode while marking UMI duplicates (in-house script and HTSeq-count). Differential expression testing was performed with DESeq2^103^ (v.1.34), and pairwise comparison was performed with lfcShrink function with -type ashr^106^. Genes with log2foldchange ≥ 1 and ≤ -1 with padj ≤ 0.05 and baseMean ≥ 10 were considered differentially expressed. Clustering was performed with the kmeans function in R.

### Splicing efficiency and alternative splicing analysis

Splicing efficiency analysis was performed as described earlier^107^. To estimate splicing efficiency at the gene level, we used a unique and non-overlapping set of introns for each gene, confidently supported by the entire RNA-seq dataset. We used only splice-site junction spanning reads for quantification and defined splicing efficiency as the ratio between exon-exon reads and all reads (exon-exon plus exon-intron) mapped to junctions. Splicing efficiency values for transcripts and introns were compared using a two-sided t-test. eCLIP clusters (peak regions) were intersected with introns using bedtools^108^. Differential splicing analysis was done using rMATS (v4.0.2)^109^ and implementing the MAJIQ-VOILA (v.2.4) pipelines ^110, 111^. Pairwise rMATS differential alternative splicing events were obtained by options -b1, -b2, -gtf, -t paired -- readLength 125 --variable-read-length. For MAJIQ analysis, confounding variations associated with the RNAseq batch were modeled and fitted with MOCCASIN^112^. To quantify local splicing variations (LSVs) and to define splice graphs, the MAJIQ build tool was used, followed by deltapsi and heterogen with default parameters. Results from deltapsi output were further analyzed and parsed with VOILA modulizer and tsv tools. P-values from heterogen output were used for the event level splice type-specific analysis. Splice graphs were visualized using the VOILA view function, and mis-spliced transcripts between comparisons were defined as significant with ΔPSI (deltaPSI) ≥ 0.2 and ≤ -0.2, p ≤0.05. For both algorithms, the gencode.vM25.annotation.gtf transcriptome was used as an input.

### Small RNA-seq library preparation

Small RNA libraries were constructed with a total of 1μg of purified RNA (as described above) using TruSeq Small RNA Library Prep or Lexogen Small RNA-seq Library Prep as per the manufacturer’s instructions with 14-16 cycles of PCR. Size selection of amplified libraries was made by running on 3% agarose gel followed by gel extraction. The quality of libraries was assessed with Agilent TapeStation before pooling. Truseq small RNA libraries were sequenced to obtain 50 bp single-end reads, and Lexogen small RNA libraries were obtained to get 75bp paired-end reads in the Illumina NextSeq 550.

### Small RNA-seq analysis

Adaptor sequences from small RNA-seq libraries were trimmed with cutadapt. Trimmed reads were aligned to the GRCm38/mm10 reference genome using STAR (version v2.4.2a) with parameters --alignEndsType EndToEnd --outFilterMismatchNmax 3 -- outFilterMultimapScoreRange 0 --outFilterScoreMinOverLread 0 -- outFilterMatchNminOverLread 0 --alignSJDBoverhangMin 1000 --alignIntronMax 1. A custom gtf file was constructed with all mature miRNA annotations from miRbase (v.22) systematically included as distinct transcripts for each parent pri-miRNA gene annotation in gencode.vM25.annotation.gtf to create a txdb object. Primary alignments were counted using the qCount function from the QuasR Bioconductor package (v.1.34) with default parameters. The batch effect was corrected using the ComBat_seq^102^ function from the sva Bioconductor package (v.3.42). Normalization and testing for differential expression on corrected read counts were performed with DESeq2^103^ on iteratively estimated size factors and mean dispersion estimates with nbinomWaldtest. Raw p-values were adjusted for multiple testing using the Independent Hypothesis Weighting ^113^ procedure from Bioconductor package IHW. Fold change was estimated manually with the following formula log2[(Nx+1)/(Ny+1)], where x and y are two conditions and N is the mean of normalized counts. Only mature miRNA with log2FC ≥ 1 and ≤ -1 with padj ≤ 0.05 and baseMean ≥ 5 were considered differentially expressed unless specified otherwise. Predictions of miRNA target genes were downloaded from miRDB (v6.0)^86^. Results were filtered to include only miRNA-mRNA pairs with opposing log2FC values in our RNAseq experiments and with miRDB score ≥ 60.

### ATAC-seq library preparation

ATAC-seq was performed on *Lis1* WT, F/-, and OE cells grown in 2i+LIF, FBS+LIF, and FGF+Activin media [supplementary Table 1] for five passages, as described earlier^114^.

Briefly, cells were treated with Trypsin and washed in ice-cold PBS. 50,000 cells were lysed with (10 mM Tris-HCl, pH 7.4, 10 mM NaCl, 3 mM MgCl2, and 0.1% IGEPAL CA-630) ice-cold lysis buffer, and nuclei were spun at 500g for 10 min in a refrigerated centrifuge.

Immediately, the pellet was resuspended and incubated in the transposase reaction mix (25 μl 2× TD buffer, 2.5 μl transposase (Illumina), and 22.5 μl nuclease-free water). The reaction was carried out at 37°C for 30 min. The DNA was directly purified using the MinElute PCR Purification kit (QIAGEN). After purification, the eluted DNA was amplified for 11-12 cycles with Kappa HiFi Ready-mix (Roche) and custom Nextera PCR primers. The libraries were cleaned up again with a MinELute PCR purification kit and were sequenced.

### ATAC-seq analysis

Nextera adaptors were removed using cutadapt, and bam files were generated by mapping trimmed reads to the genome with Bowtie2 parameters –sensitive local -k 4 and the read pairs separated by more than 2kb were not considered. Further, post mapping alignments were filtered by removing reads mapped to chrM, PCR duplicates with PICARD markdup, and multimapped reads were removed with samtools -q (MAPQ<30)^115^. To identify chromatin accessible regions, only short reads (≤ 130bp) that correspond to the nucleosome-free region were considered. The TOBIAS framework was implemented for subsequent ATAC-seq analysis^116^. Briefly, ATAC-seq peaks were identified by running MACS2 with parameters -- nomodel --shift -100 --extsize 200 --keep-dup all -q 0.01 for each processed bam file. Further, processed bam files for each treatment and condition were merged, and Tn5 transposase insertion bias was corrected with ATACorrect. Footprint scores were generated around merged narrow peak file output from peak calling with ScoreBigwig tools after removing Blacklisted regions [https://www.encodeproject.org/files/ENCFF547MET/@@download/ENCFF547MET.bed.gz]. Transcription factor motifs were obtained from HocomocoV11, JASPAR Core 2020, and the JASPAR PBM Homeo collections. Redundant motifs between databases were filtered to one motif for each transcription factor available for the mouse and mouse-human conserved motifs to obtain 645 motifs in JASPAR format. Bias corrected footprint scores were compared to predict transcription factor binding scores using the BINDetect tool, and a comparison was made between conditions for each treatment and across treatments. Primary alignments were quantified on bias-corrected peak regions using qCount from the QuasR Bioconductor package (v.1.34).

Batch correction was performed with Combat_seq and design matrix ∼Treatment+Condition: Treatment was used. Differentially accessible regions on corrected primary alignments were obtained with DESeq2^103^ Bioconductor package (v.1.34) using nbinomWaldTest (betaPrior = TRUE). Pairwise contrasts were obtained by creating a model with group design for condition comparison across each treatment, and raw p-values were adjusted for multiple testing using Benjamini and Hochberg procedure^117^. Open chromatin regions with log2foldchange ≥ 1 and ≤ - 1 with padj ≤ 0.05 and baseMean ≥ 5 for condition comparison in each treatment and across treatments were considered differentially accessible (n=6984).

### seCLIP-seq

seCLIP libraries were generated based on the standardized experimental protocol as previously reported^118^ with slight modifications. Briefly, V6.5 mESCs were expanded in a 15 cm plate in FBS+LIF media to a density of 30x10^6^ cells. Cells were washed on ice and collected in cold 1X PBS to UV cross-link at 400 mJ/cm^2^. Sonication was performed for 2 min in a Covaris E220 instrument at intensity 140, burst 200, and duty 5. Lysates were treated with RNase I (1:25 dilution; Ambion) for 3 min at 37°C, and the clarified lysate was transferred to protein G Dynabeads (Invitrogen) conjugated to the monoclonal anti-LIS1 antibodies for two biological replicates. On beads, dephosphorylation was performed with FastAP (ThermoScientific). A 3’ RNA adapter /5’Phos/rArGrArUrCrGrGrArArGrArGrCrArCrArCrGrUrC/3’SpC3 was ligated to the samples using T4 RNA Ligase (NEB, M0437M) at room temperature for 75 min.

Phosphorylation was performed on bead using PNK (NEB, M0201L) and washed once with wash buffer (20 mM Tris-HCl pH 7.4 ,10 mM MgCl2,0.2% Tween-20), once with high salt buffer (50 mM Tris-HCl pH 7.4 ,1 M NaCl ,1 mM EDTA, 1% NP-40, 0.1% SDS, 0.5% sodium deoxycholate) and then twice with wash buffer. Samples were eluted by heating to 70°C for 10 minutes in 1X NuPAGE LDS loading buffer and 100mM DTT at 1,200rpm. Eluates were resolved by denaturing gel electrophoresis and transferred onto a nitrocellulose membrane to determine the migration of Protein-RNA complexes. The relevant region of the separated complexes was cut into small pieces on Whatman paper and transferred to Eppendorf tubes for library preparation. The RNA from the membrane was then isolated by digesting with proteinase K solution [32 units of proteinase K (Invitrogen), 100mM Tris pH 7.5, 50mM NaCl, 1mM EDTA, 0.2% SDS] and incubating at 50°C for 1 hour at 1,200rpm. RNA was purified by phenol-chloroform extraction followed by ethanol precipitation. The precipitated RNA from the aqueous layer was reverse transcribed using Superscript III reverse transcriptase (Invitrogen) and primer (5’- CAGACGTGTGCTCTTCCGA-3’). A 3′ DNA linker /5’Phos/NNNNNNNNNNAGATCGGAAGAGCGTCGTGT/3’SpC3 was ligated onto the cDNA product with T4 RNA ligase. Libraries were amplified with the TruSeq LT adapters, and PCR products were gel-purified using a 3% agarose gel^119^.

### seCLIP-seq analysis

seCLIP clusters enriched against size-matched input (SMI) were identified as described previously^120^. Sequenced reads were processed to remove inline barcodes and adaptor sequences. Trimmed reads mapped to a repeat element database (RepBase v25.06) were removed, and unmapped reads were then mapped to mouse mm10 reference genome assembly using STAR (v2.7.7). Aligned deduplicated reads were merged, and peaks were called using Clipper.

Normalized peak files were ranked by entropy score as inputs to IDR to determine reproducible peaks. Reproducible peaks were annotated based on overlap with gencode.vM25.annotation.gtf transcripts. Motif enrichment analysis was performed using the 50 nt sequences flanking the center of seCLIP clusters in both directions. The background sequence sets were generated by HOMER (v4.11)^121^ for all possible known motifs. The ngs.plot.r^122^ package was used for metaplots. For the eCLIP map of MAJIQ events, coordinates of cassette exons with retained introns and median ΔPSI difference of 0.2 between conditions were considered. These regions were extended to the intronic region 250 nt upstream of the 3’ splice site plus 50 nt downstream and 50 nt upstream of the 5’ splice site plus the intronic region 250 nt downstream. The seCLIP reads were modeled on the cassette exon extended regions with qProfile functions from the QuasR Bioconductor package (v.1.26.0)^101^. Metaprofiles were normalized to reads per kilobase per million over SMI and averaged over 50 bp for visualization.

For LIS1 seCLIP and miRNA expression joint analysis, intron coordinates for protein-coding genes from gencode.vM25.annotation.gtf were extracted using the extract_pc and extract_introns function from the GencoDymo R package https://doi.org/10.5281/zenodo.3605996. All the non-redundant introns and only one assignment from redundant introns were included in the analysis. The remaining introns other than protein-coding from gencode.vM25.annotation.gtf were categorized as non-protein-coding introns. All the mature miRNAs from miRbase(v22) showing overlap within 2000 nt distance upstream and downstream from LIS1 seCLIP peaks (<2kb) were extracted with bed_closest function using valr (v0.6.4)^123^ R package. miRNA coordinates overlapping with a distance more than 2000 nt upstream and downstream of LIS1 seCLIP peaks and within introns of protein-coding (>2kb protein-coding) as well as non-protein-coding (>2kb non-protein coding) genes were obtained using bed_closest from valr package. Log2 of baseMean values of DESeq2 normalized miRNA counts from LIS1-OE, WT, and F/- for the above three categories were compared along with the remaining miRNAs (categorized as other miRNAs). For the LIS1 seCLIP miRNA profile, pre-miRNA coordinates were extended upstream and downstream 2000nt, and the matrix was built using deepTools (v3.5.1)^124^. Defined regions were scaled up to 5000 nt, and binning was done based on the median value.

### AFM experiment and analysis

Atomic force microscopy (AFM) experiments were carried out on TT-FHAgo2 and TT- FHAgo2 LIS1-OE mESCs with and without doxycycline-induced expression of AGO2. Cells were cultured on 5-cm-diameter tissue culture dishes coated with Matrigel (Corning Life Sciences) and grown in Serum+LIF conditions. Cells were allowed to grow for 3 days to reach confluency without colonies touching each other. Fresh media was added before the AFM measurements. AFM imaging and stiffness maps were measured on a Nanowizard III AFM (JPK/Bruker, Berlin, Germany) in QI mode operated with the JPK control software v.6.1.159. In this mode, force-distance curves are collected at each pixel and used to generate topographic images simultaneously with nano-mechanical data. A BioAC-CI CB3 probe (Nanosensors) was used (nominal spring constant ≈ 0.06 N/m), and sensitivity and spring constant were calibrated before each measurement using the non-contact calibration procedure in the JPK software. The maximum force was chosen to keep penetration depths between 150-500nm. The elastic modulus maps were calculated by applying a Hertzian model (using JPK data processing software v6.1.86), presuming a Poisson ratio of 0.5 and conical tip shape with 22° half-cone angle. The modulus values were extracted from the modulus maps for the regions in the centers of the colonies, defined by the corresponding height images using Gwyddion software (v2.60)^125^.

### Quantification and statistical analysis

All bedgraph files were prepared using deepTools (v3.5.1). seCLIP bedgraph were RPM normalized. RNA-seq, small RNA-seq and ATACseq bedgraphs were normalized with scaleFactors. For calculating scaleFactors, reciprocals of DESeq2 size factors were estimated using calcNormFactors from edgeR Bioconductor package with RLE method. Track plots were generated from bedgraph files with SparK^126^.

All the statistics were performed in base R or using rstatix and statExpressions packages^127^ R packages. Gene set over-representation test (p.adj <0.05) and GO term network was done using the Bioconductor package clusterprofiler (v4.2.1)^128^. Schematics related to immunoprecipitation and mass spectrometry were created with Biorender [https://biorender.com/].

### miRNA qRT-PCR

The cDNA for qRT experiments was prepared with miScript II RT kit (Qiagen, 218161) using HiFlex buffer. The qRT reactions were performed with miScript SYBR Green PCR kit (Qiagen, 218075). The values of miRNAs were normalized to that of U6 snRNA. The following primers were used for the reactions:

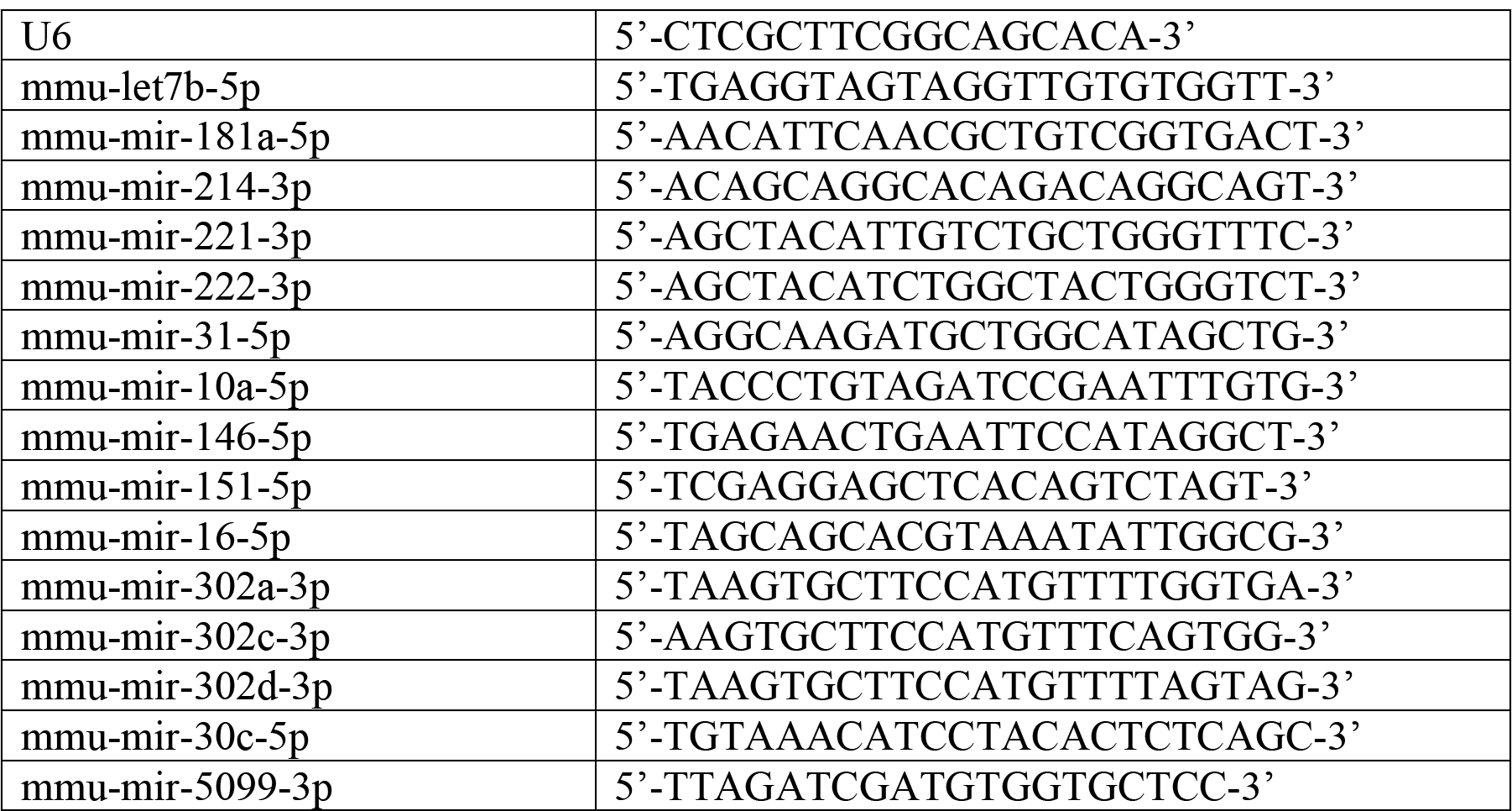
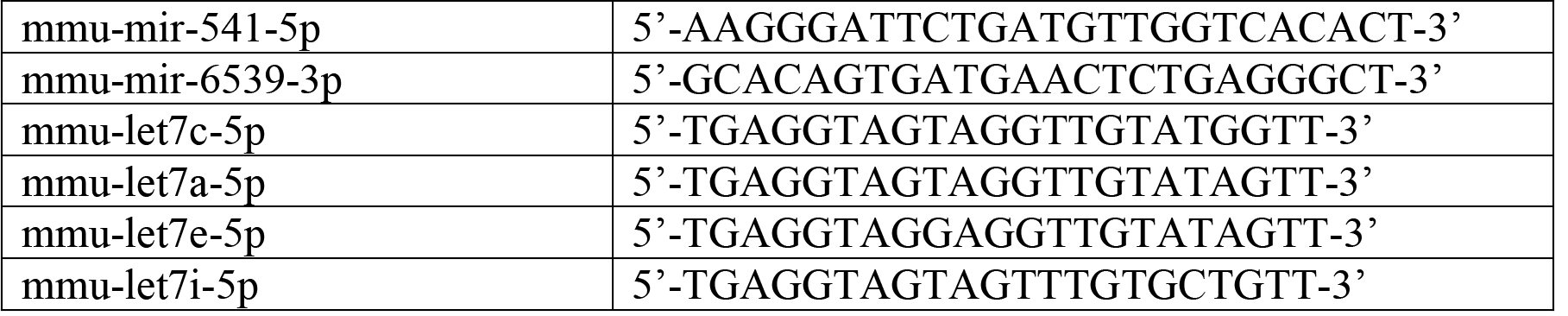

The reverse primer was miScript Universal Primer (Qiagen).

### Genotyping

Mice were genotyped at 21 days old by established methods using following primers: 129S-Pafah1b1^tm2Awb^/J- (Lis1 *^Flox/Flox^*, Lis1 *^Flox/WT^* or LIS1 *^Flox/-^*): Lis1 wild-type and floxed alleles were detected with

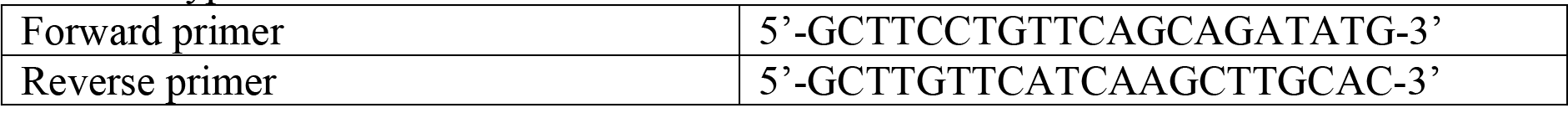

 and deleted allele was detected with

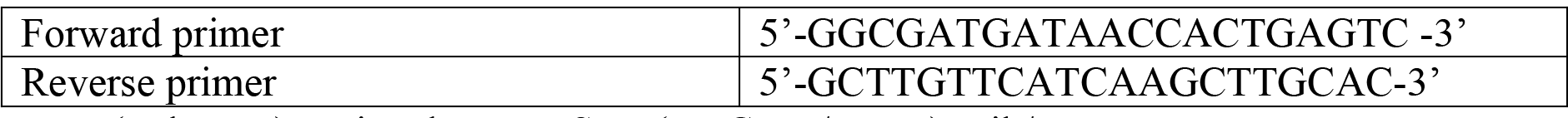

 B6-Tg(Pgk1-cre)1Lni and B6;129S-Tg(UBC-cre/ERT2)1Ejb/J: Cre transgene was detected with

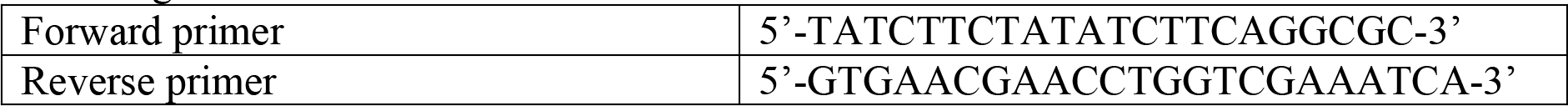

129S;ICR-*Tg(CAGG-loxP-LacZ-neo-loxP-PAFAH1B1-DsRed*):
Lis1-DsRed transgene was detected with

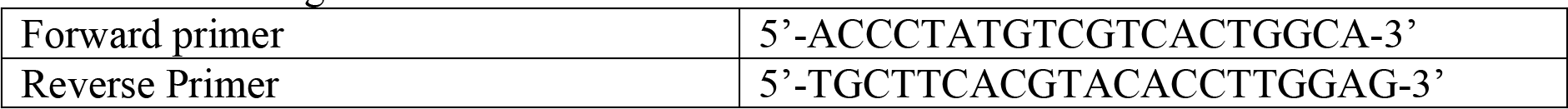

## Extended data and Methods

**Extended Data Fig. 1:**
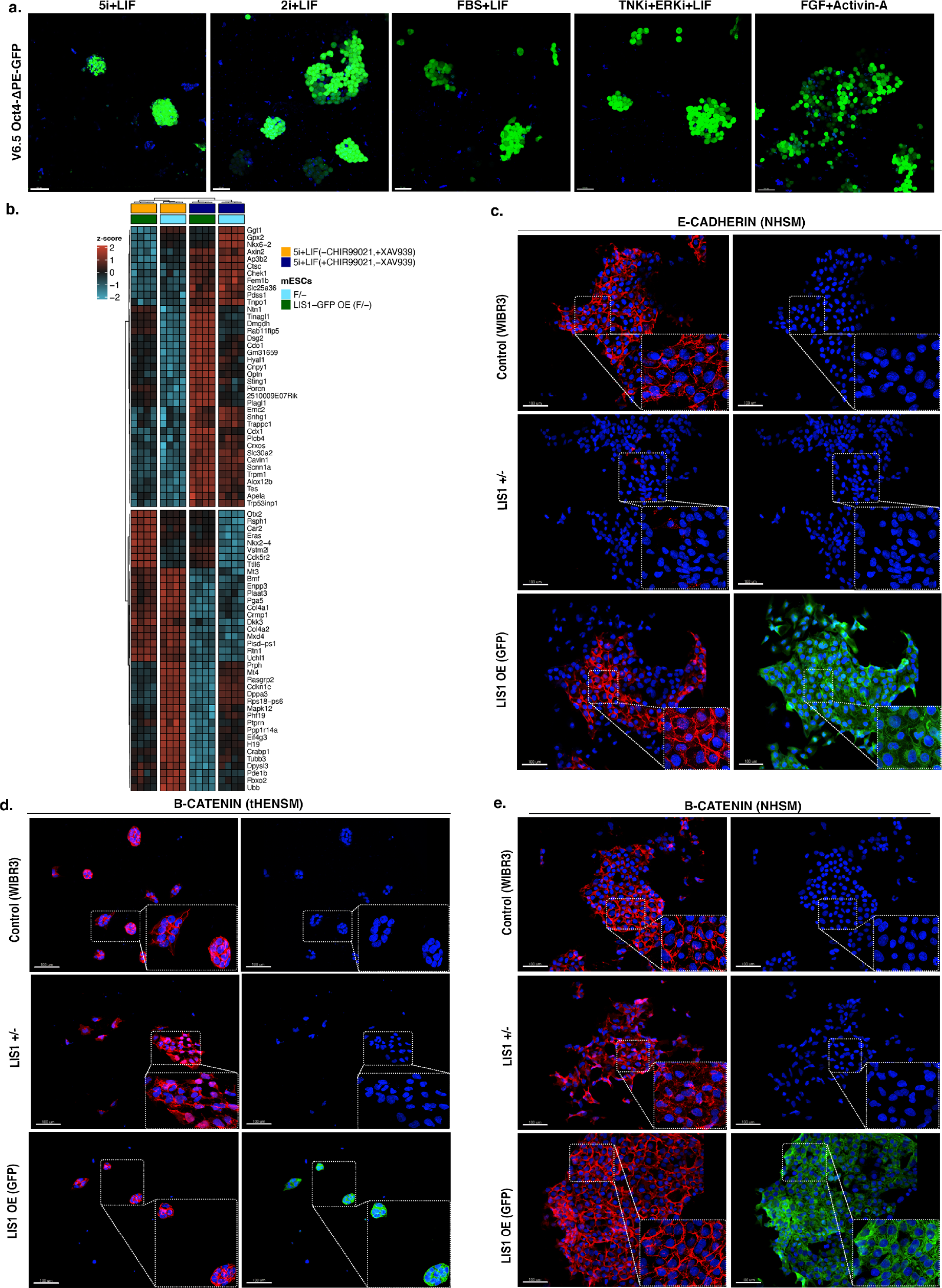
WNT signaling influences sustenance of mutant LIS1 embryonic stem cells. **a.** Fluorescent images of V6.5 Oct4-ΔPE-GFP mESCs cultured using different media. Left to right: 5i+LIF, 2i+LIF, FBS+LIF, Tnki+Erki+LIF and FGF+Activin. **b.** A heatmap showing LIS1 dose-dependent gene expression changes common between F/- and LIS1-GFP- OE(F/-) mESC isogenic lines for comparison in 5i+LIF media with WNT activator (CHIR99021) and WNT inhibitor (XAV939). The data are shown on a Z-score scale of the variance stabilizing transformation on normalized reads. **c-e.** Representative images of immunostainings of WIBR3 (Control), LIS1+/- and pB-LIS1GFP overexpression isogenic hESCs **c.** hESCs cultured in NHSM media and immunostained with anti-E-Cadherin antibodies. **d.** and **e.** Immunostainings of hESCs cultured in tHENSM and NHSM media, respectively, using anti-β-Catenin antibodies (scale bars, 100 μm. Insets represent 2.5x zoom).

**Extended Data Fig. 2:**
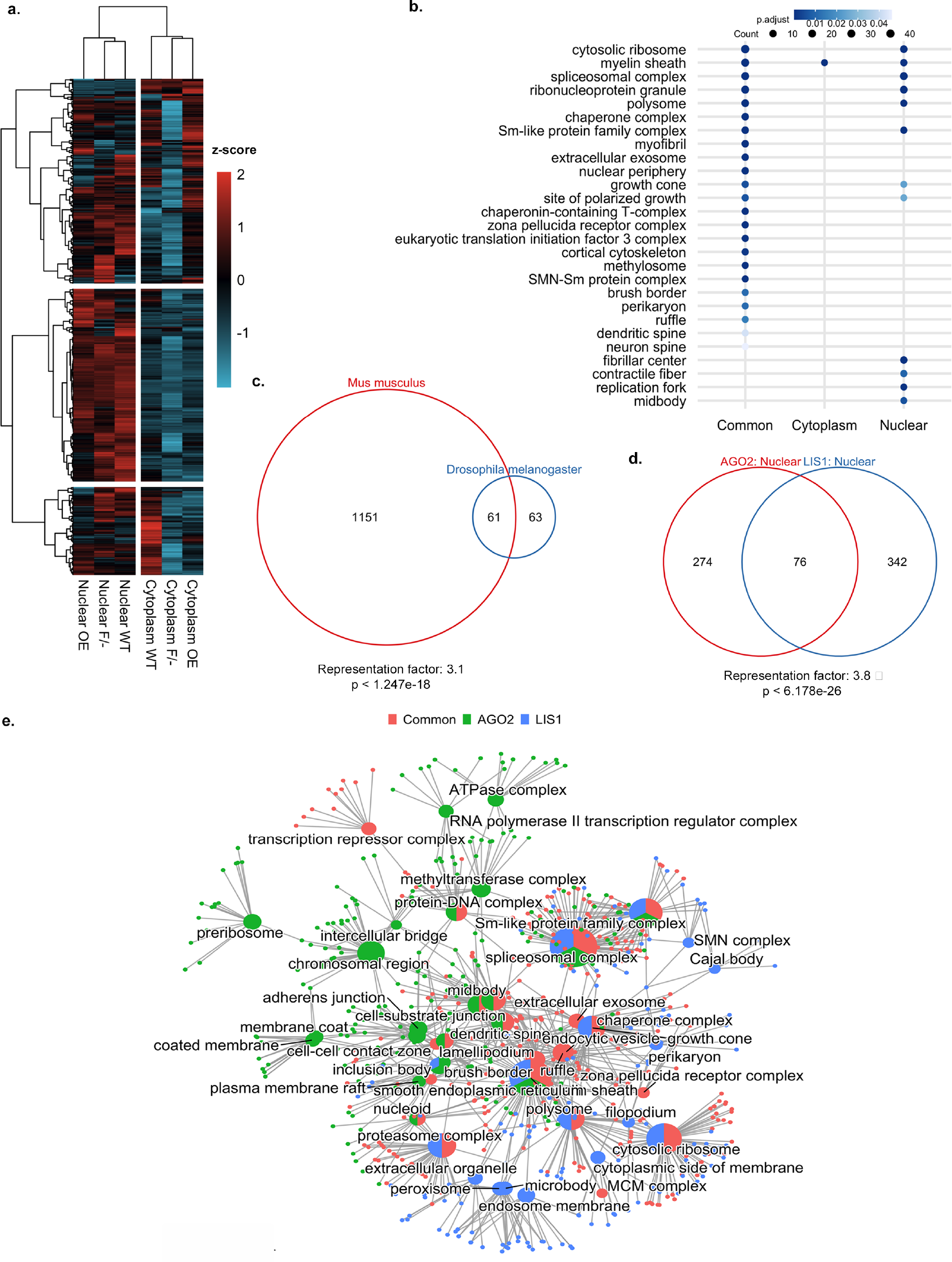
The LIS1 interactome. **a.** A heatmap of normalized peptide intensity on a log2 scale of proteins immunoprecipitated with anti-LIS1 antibodies using nuclear and cytoplasmic fractions from LIS1-dsRED overexpression, WT, or F/, identified by mass spectrometry, after scaling. **b.** Overrepresentation test analysis for GO terms of LIS1 interacting proteins. Categorized as common for proteins found in nuclear and cytoplasmic fractions, nuclear only, or cytoplasmic only. **c.** A Venn diagram showing the overlap of LIS1 interacting proteins identified in this study (all fractions/extracts) and *Drosophila melanogaster* orthologs from Guruharsha *et al*.^50^. The significance of the overlap was tested with the hypergeometric test. **d.** A Venn diagram showing the overlap of LIS1 interacting proteins identified in nuclear extracts of LIS1-dsRED OE, WT, and F/- with AGO2 nuclear interacting proteins from Sharshad *et al.* ^51^. The significance of the overlap was tested with the hypergeometric test. **e.** The network of GO terms showing linkage for LIS1 interacting proteins from this study and AGO2 interacting proteins from Sharshad *et al.* ^51^. Big hub nodes with pie represent GO terms for the distribution of proteins specific to LIS1 (blue), AGO2 (green), and common (red) interacting proteins between the two. Pie size corresponds to the total number of proteins in each GO term. The small nodes represent proteins linked to GO term nodes.

**Extended Data Fig. 3:**
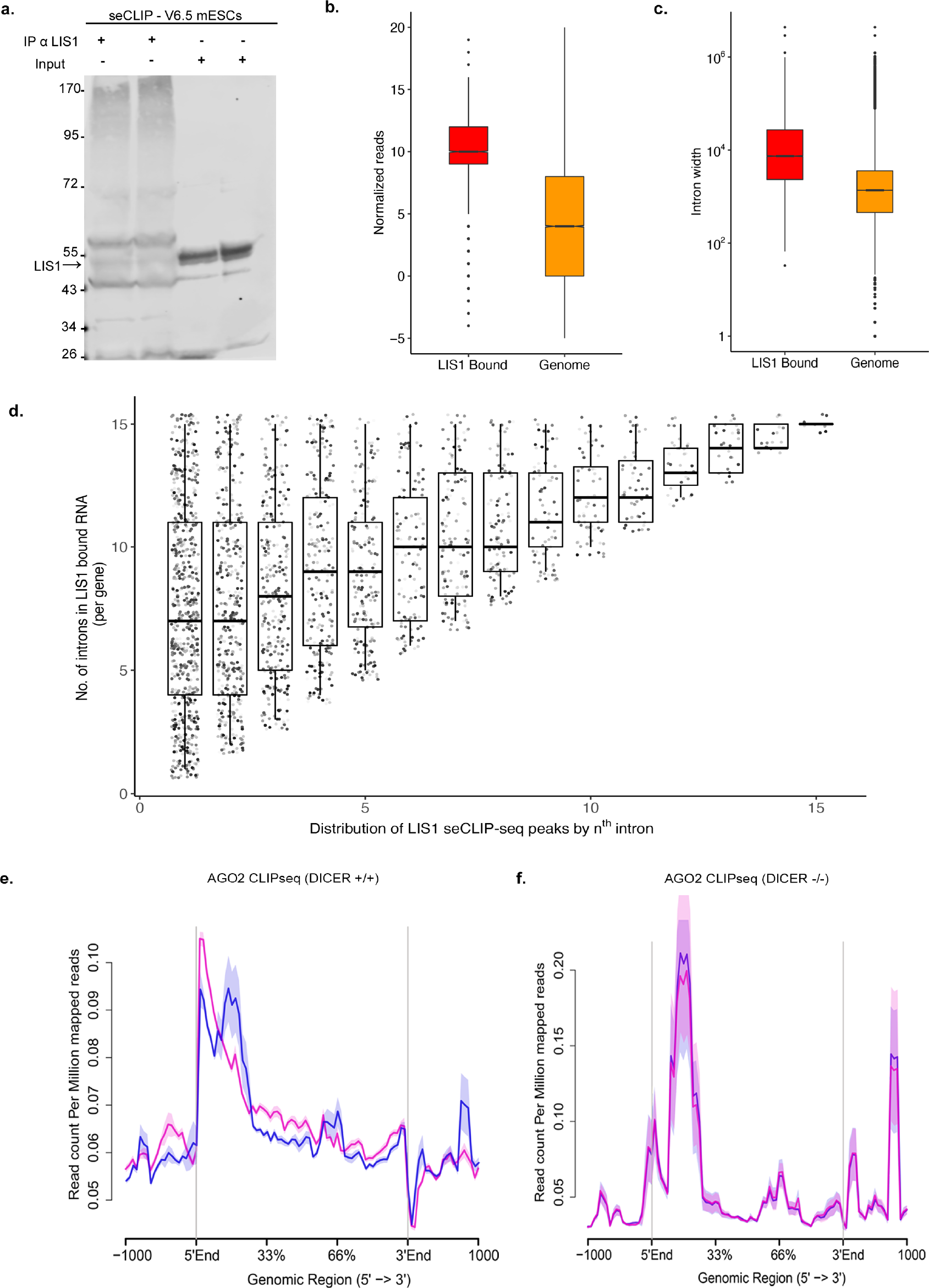
Combinatorial regulation of pre-mRNA processing by LIS1. **a.** A western blot of LIS1 immunoprecipitation from V6.5 mESCs in the seCLIP experiment (two biological replicates). **b.** The expression level of LIS1 bound genes compared to the whole genome. Values are log2 (baseMean) as calculated by DESeq2. **c.** A boxplot of LIS1 bounded intron lengths versus the entire genome introns (Gencode vM25). **d.** LIS1 seCLIP-seq clusters as a function of intron number within a gene (x-axis) and the total number of introns (1-15 introns) in LIS1 bound genes (y-axis). **e.** and **f.** Metagene plot of AGO2 CLIP-seq data from the study of Leung *et al.* ^56^ in (**e**) WT embryonic stem cells DICER(+/+) and (**f**) DICER knockout embryonic stem cells. Statistics for 3b,c are in Extended Data Table 7.

**Extended Data Fig. 4:**
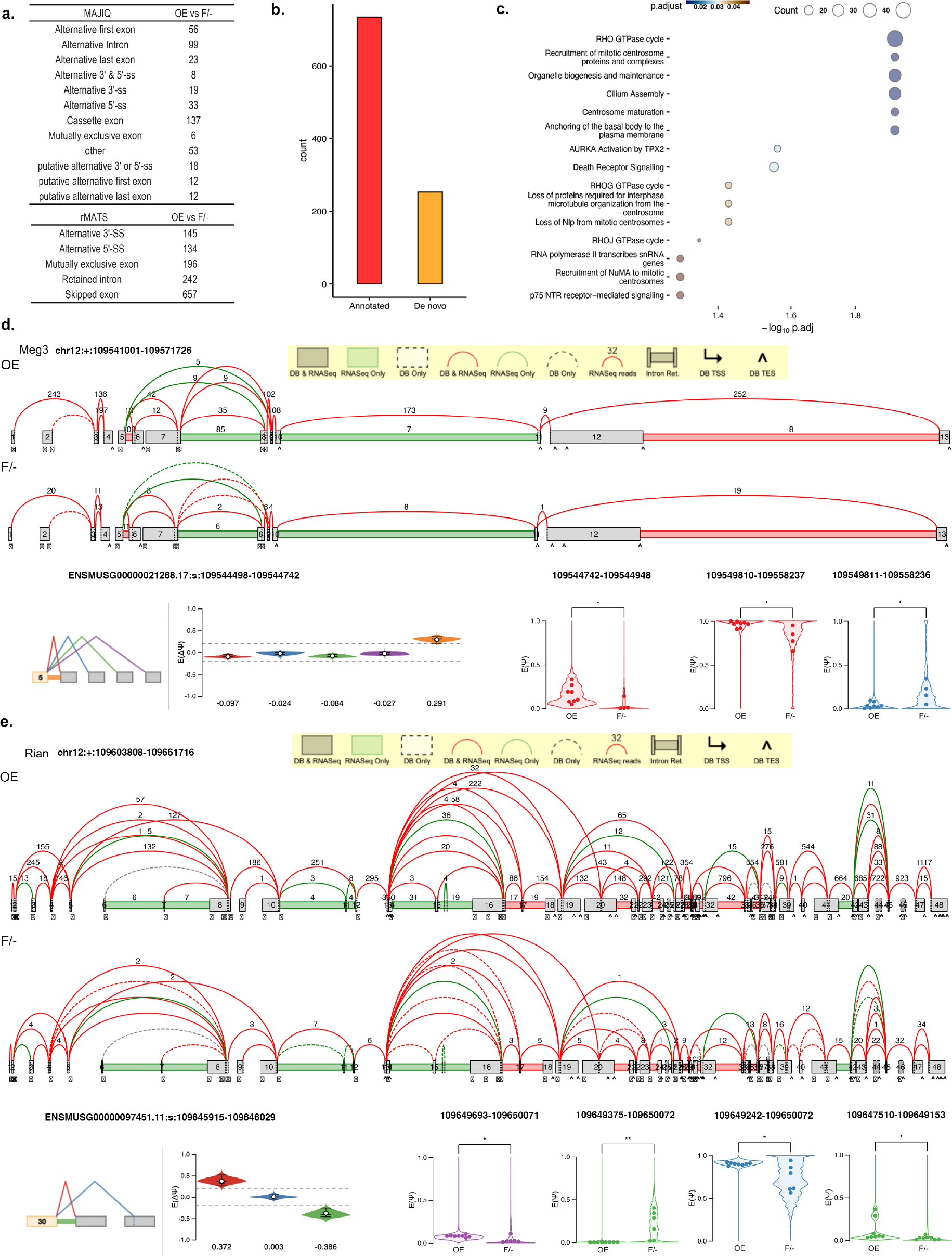
LIS1 dosage affects RNA splicing. **a.** A table of all differential alternative splicing events between the LIS1 OE and F/- genotypes identified with MAJIQ and RMATS. **b.** The number of annotated versus *de novo* events identified by MAJIQ**. c.** Pathway enrichment analysis of the genes that harbor differentially spliced events. The analysis was performed with the genes that were found in both MAJIQ and rMATS analysis, using the Reactome database. **d.** and **e.** Splice graphs on (top) and representative significant splice events (bottom) of the two long non-coding RNAs *Meg3* and *Rian* from the *Meg3-Mirg* locus in the mouse.

**Extended Data Fig. 5:**
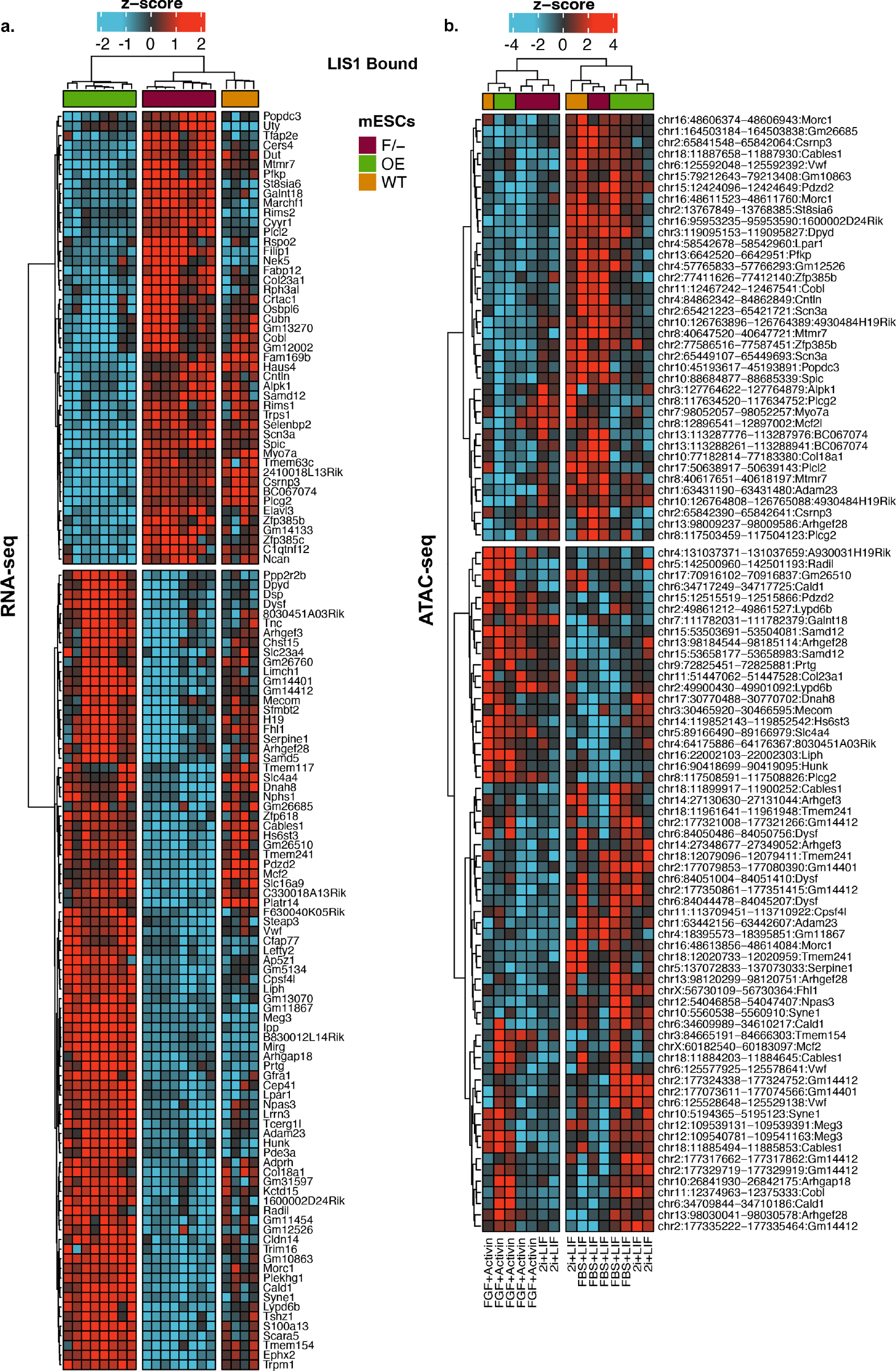
Gene expression and chromatin accessibility changes of LIS1 bound genes. **a.** A heatmap showing the transcriptional changes between OE and F/- with WT mESCs in FBS+LIF for 134 LIS1 bound in seCLIP-seq and differentially expressed genes in RNAseq data. The data are shown on a Z-score scale of the variance stabilizing transformation on normalized reads. **b.** A heatmap showing differential accessibility of 79 open chromatin regions associated with differentially expressed and LIS1 bound genes. The comparison is shown for OE and F/- transition across naïve and primed states (2i+LIF, FBS+LIF, and FGF+Activin). The data are displayed on a Z-score scale of the variance stabilizing transformation on normalized primary alignment counts.

**Extended Data Fig. 6:**
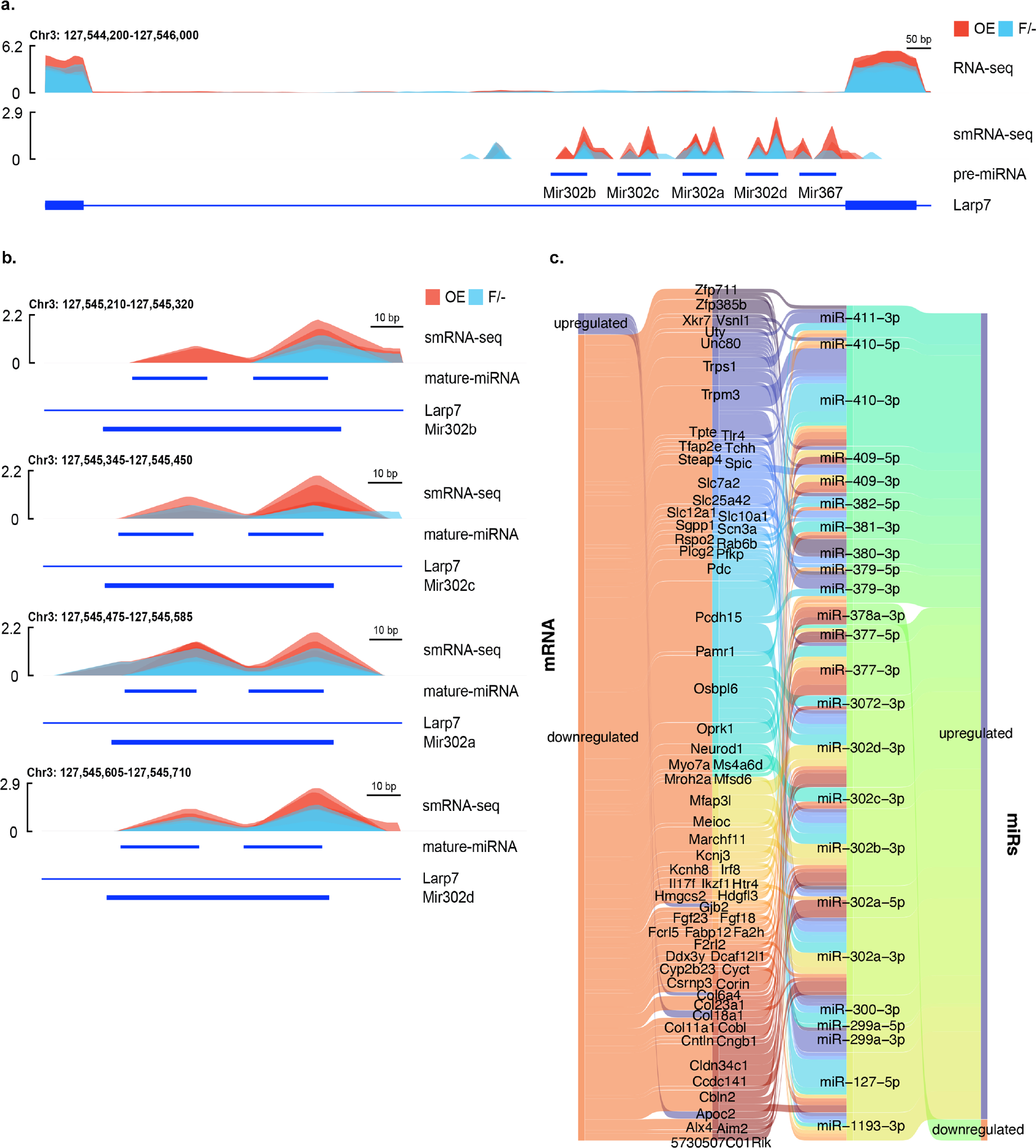
LIS1 affects the expression of miRs and target genes. **a.** Read coverage of small RNA-seq and total RNA-seq in the miR302-367 cluster (OE in red and F/- in blue). **b.** small RNA-seq read coverage of miR302b, miR302c, miR302a, and miR302d (top to bottom). OE in red and F/- in blue **c.** Sankey diagram showing miRNA target analysis. Right: differentially expressed miRNAs between OE and F/- conditions. Left: differentially expressed genes between the same conditions. The identification of the miRNA targets was done using the miRDB database^86^, only miRNA-RNA pairs with opposite expression change were selected.

**Extended Data Fig. 7:**
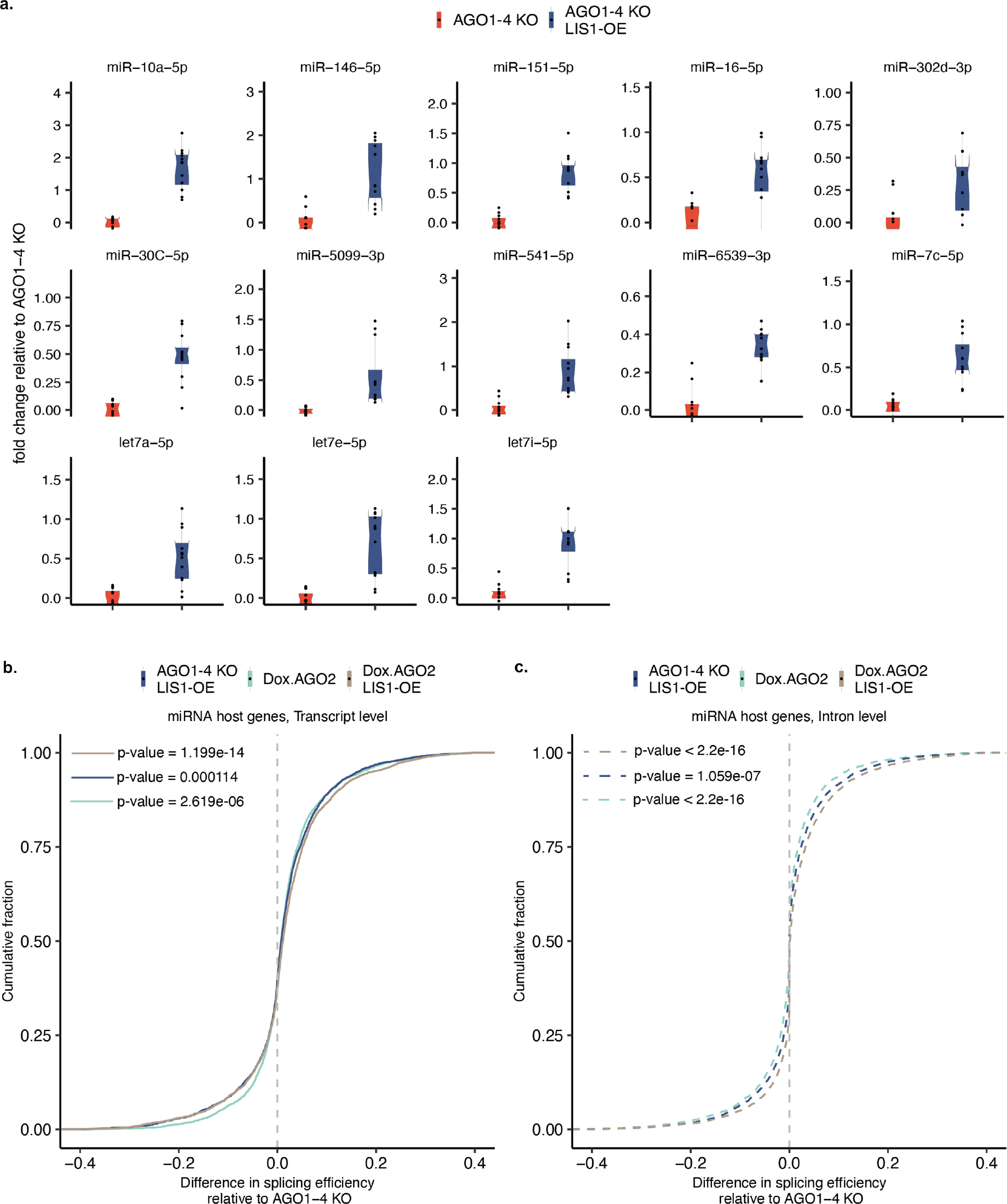
LIS1 OE affects expression miRs and processing of miR host transcripts in AGO1-4 KO mESCs. **a.** qRT-PCR validation of a subset of mature miRs that were found to be upregulated by overexpression of LIS1 on the background of AGO1-4 KO. (n = 4 (x3), all p-values are reported in Extended Data Table 7). **b.** Intron level quantification of differences in splicing efficiency for Dox. AGO2 ^AGO1-4KO^, LIS1-OE ^AGO1-4KO,^ and Dox. AGO2 LIS1-OE ^AGO1-4KO^ relative to AGO1-4KO in miRNA host genes. **c.** Transcript level quantification of differences in splicing efficiency for Dox. AGO2 ^AGO1-4KO^, LIS1-OE ^AGO1-4KO,^ and Dox. AGO2 LIS1-OE ^AGO1-4KO^ relative to AGO1-4KO in miRNA host genes.

**Extended Data Fig. 8:**
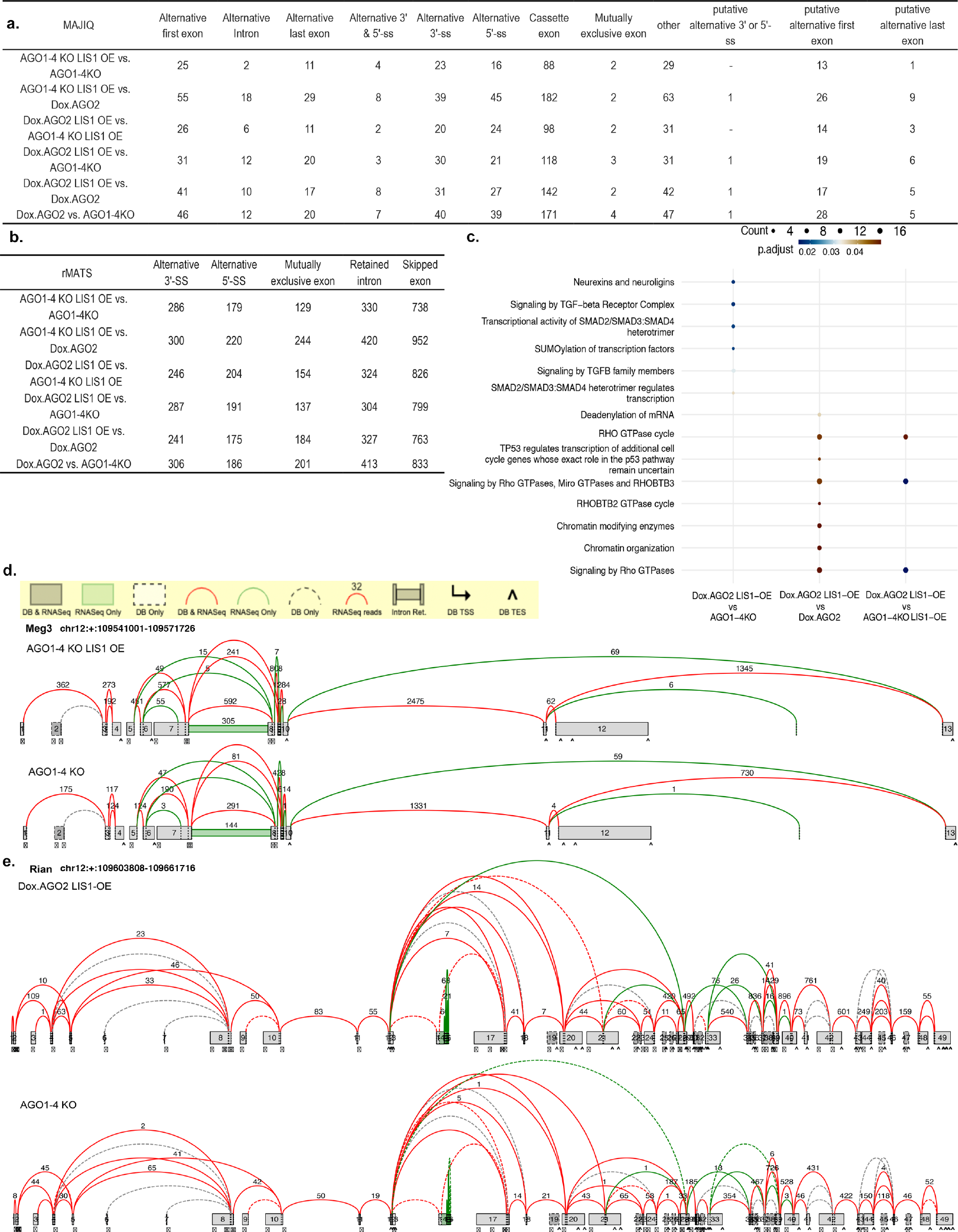
LIS1 and AGO2 effects on RNA splicing. a-b. A table of all differential alternative splicing events between the AGO1-4 KO, Dox.AGO2 ^AGO1-4KO^, LIS1-OE ^AGO1-4KO^, and Dox.AGO2 LIS1-OE ^AGO1-4KO^ genotypes identified with MAJIQ and RMATS **c.** Pathway enrichment analysis of the genes that harbor differentially spliced events. The analysis was performed with the genes that were found in both MAJIQ and rMATS analysis, using the Reactome database for the comparison between Dox.AGO2 LIS1-OE ^AGO1-4KO^ and AGO1-4 KO, Dox.AGO2 LIS1-OE ^AGO1-4KO^ and Dox.AGO2 ^AGO1-4KO^, and Dox.AGO2 LIS1-OE ^AGO1-4KO^ and LIS1-OE ^AGO1-4KO^. **d.** Splice graphs for *Meg3* comparing LIS1-OE ^AGO1-4KO^ and AGO1-4 KO. **e.** Splice graphs for *Rian* comparing Dox.AGO2 LIS1-OE ^AGO1-4KO^ and AGO1-4 KO.

**Extended Data Fig. 9:**
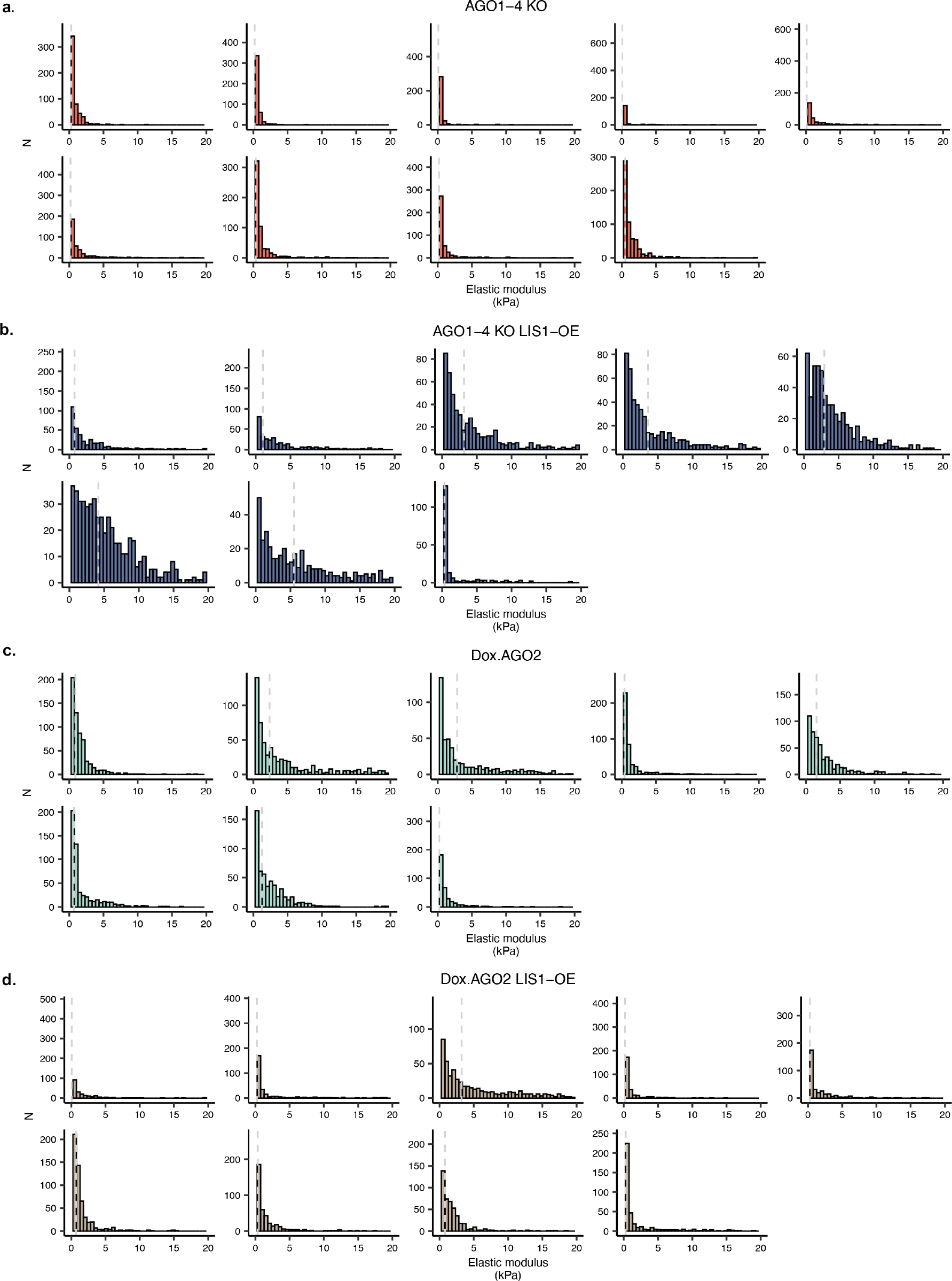
LIS1 overexpression affects the elastic modulus of AGO1-4 KO mESCs. Elastic modulus histograms for embryonic stem-cell colonies. **a.** AGO1-4 KO (n=9). **b.** LIS1-OE ^AGO1-4KO^ (n=8). **c.** Dox.AGO2 ^AGO1-4KO^ (n=8). **d.** Dox.AGO2 LIS1-OE ^AGO1-4KO^ (n=8).

